# Translation and codon usage regulate Argonaute slicer activity to trigger small RNA biogenesis

**DOI:** 10.1101/2020.09.04.282863

**Authors:** Meetali Singh, Eric Cornes, Blaise Li, Piergiuseppe Quarato, Loan Bourdon, Florent Dingli, Damarys Loew, Simone Proccacia, Germano Cecere

## Abstract

In the *Caenorhabditis elegans* germline, thousands of mRNAs are concomitantly expressed with antisense 22G-RNAs, which are loaded into the Argonaute CSR-1. Despite their essential functions for animal fertility and embryonic development, how CSR-1 22G-RNAs are produced remains unknown. Here, we show that CSR-1 slicer activity is primarily involved in triggering the synthesis of small RNAs on the coding sequences of germline mRNAs and post-transcriptionally regulates a fraction of targets. CSR-1-cleaved mRNAs prime the RNA-dependent RNA polymerase, EGO-1, to synthesize 22G-RNAs in phase with ribosome translation in the cytoplasm, in contrast to other 22G-RNAs mostly synthesized in germ granules. Moreover, codon optimality and efficient translation antagonize CSR-1 slicing and 22G-RNAs biogenesis. We propose that codon usage differences encoded into mRNA sequences might be a conserved strategy in eukaryotes to regulate small RNA biogenesis and Argonaute targeting.

## INTRODUCTION

In animals, small RNAs that are expressed in the germline and transmitted to embryo act as a defense mechanism to repress foreign RNAs such as viruses, transposons and other repetitive elements (REs). These small RNAs are essential for fertility and genome integrity (Ghildiyal and Zamore, 2009; Okamura and Lai, 2008). Their function is controlled by the conserved family of Argonaute proteins (AGOs), which loads the small RNAs and function to repress complementary mRNA targets through their endonuclease activity or by recruiting other effector silencing proteins (Grishok et al., 2001; Hammond et al., 2000; Tabara et al., 1999; Williams and Rubin, 2002). The *C. elegans* germline contains a complex small RNA regulatory network, with different classes of small RNAs, multiple AGO effectors and diverse biogenesis pathways (Billi et al., 2014). One of the most abundant class of endogenous small RNAs in the germline is the 22G-RNAs, which are single-stranded antisense small RNAs produced by RNA-dependent RNA polymerase (RdRPs) as part of an amplification system to silence target transcripts (reviewed in (Billi et al., 2014)). The production of 22G-RNAs targeting REs is triggered by over 15,000 PIWI-interacting RNAs (piRNAs or 21U-RNAs) and loaded by Worm-specific Argonautes (WAGOs) to silence REs (Bagijn et al., 2012; Batista et al., 2008; Das et al., 2008; Lee et al., 2012). 22G-RNAs are also produced from the majority of germline-expressed mRNAs by the RdRP EGO-1 and loaded into the Argonaute CSR-1 (Claycomb et al., 2009; Maniar and Fire, 2011). In contrast to the 22G-RNAs antisense to REs, which can be triggered in response to piRNAs, the primary trigger for generating CSR-1 22G-RNAs and why many germline mRNAs become targeted by CSR-1 is still unknown (Figure S1).

Given that the *C. elegans* piRNAs can trigger the silencing of their targets by imperfect complementarity, and therefore can potentially target germline-expressed mRNAs (Seth et al., 2018; Shen et al., 2018; Zhang et al., 2018), the targeting by CSR-1 22G-RNAs can function as an anti-silencing mechanism to protect germline mRNAs from piRNAs silencing (Claycomb et al., 2009; Seth et al., 2013; Wedeles et al., 2013). The anti-silencing function of CSR-1 was established with single-copy transgenes (Seth et al., 2013; Shen et al., 2018; Wedeles et al., 2013). However, germline mRNAs remain protected from piRNAs silencing even in the absence of CSR-1 (Zhang et al., 2018), and sequence-encoded features of germline mRNAs have also been proposed to prevent piRNA silencing (Seth et al., 2018; Zhang et al., 2018). To what extent endogenous germline-expressed genes are regulated by the antagonistic functions of CSR-1 and piRNA pathways remains elusive (Figure S1).

In addition, CSR-1 has been proposed to directly regulate the expression of its germline targets. Of the Argonautes that load 22G-RNAs, only CSR-1 has demonstrated slicer activity on target mRNA *in vitro* (Aoki et al., 2007). Worms lacking CSR-1 protein display downregulation of germline CSR-1 targets (Cecere et al., 2014; Claycomb et al., 2009; Conine et al., 2013), those expressing a CSR-1 catalytic mutant protein show upregulation of its germline target genes (Gerson-Gurwitz et al., 2016). Therefore, the gene regulatory functions of germline CSR-1 22G-RNAs remain incompletely understood (Figure S1).

Many germline Argonautes, including CSR-1 and PIWI, and proteins involved in 22G-RNA biogenesis, including RdRPs, localize to perinuclear condensates called germ granules (Claycomb et al., 2009; Gu et al., 2009). These germ granules are thought to be the site for the biogenesis of all germline 22G-RNAs. However, disruption of specific germ granule, the mutator foci, which participates in piRNA-dependent 22G-RNA accumulation, has no apparent effect on CSR-1 22G-RNAs (Phillips et al., 2012; Zhang et al., 2011). Moreover, the type of RNA template used by the EGO-1 RdRP to generate CSR-1 22G-RNAs also remains mysterious. During exogenous RNAi, the addition of polyUG to cleaved mRNA targets by RDE-3 recruits RdRPs EGO-1 and RRF-1 to synthesize 22G-RNAs (Preston et al., 2019; Shukla et al., 2020). However, RDE-3 is not required to generate CSR-1 22G-RNAs (Gu et al., 2009; Shukla et al., 2020). Thus, whether CSR-1 22G-RNAs are synthesized in germ granules on a specific type of RNA substrates remains to be elucidated.

In the current study, we elucidate CSR-1 catalytic activity-dependent and independent germline gene regulation and decipher the rules governing CSR-1 22G-RNA biogenesis. We demonstrate that the slicer activity of CSR-1 triggers the biogenesis of 22G-RNAs on the coding sequence of germline mRNAs. We establish that CSR-1 22G-RNAs are synthesized on an actively translated mRNA template in the cytosol, independent of germ granules. Overall, this study establishes that translation and codon usage dictate CSR-1 slicer activity on a target mRNA to regulate small RNA biogenesis and functions.

## RESULTS

### Defects in CSR-1 catalytic activity mainly impact 22G-RNA abundance

Worms lacking CSR-1 show downregulation (Cecere et al., 2014; Claycomb et al., 2009; Conine et al., 2013) whereas those expressing CSR-1 protein in which the catalytic DDH motif was mutated to ADH show upregulation of some targets (Gerson-Gurwitz et al., 2016). The global impact of CSR-1 mutations on gene expression might depend on the developmental context and might be biased by developmental defects (Claycomb et al., 2009; Yigit et al., 2006). Indeed, we observed differences during oogenesis in the adult *csr-1* catalytic mutant (*csr-1* ADH) and knockout (*csr-1* KO) worms marked by a delayed onset of oocyte production and increased accumulation of oocytes in germline at more advanced stages compared to wild-type (WT) (Figure S2A-C). To overcome this limitation, we developed a sorting strategy to obtain a synchronized population of WT and first-generation homozygotes for *csr-1* KO or CSR-1 ADH strains using COPAS biosorter. Using this strategy, we enriched for larval stage late L4 worms lacking the germline developmental abnormality, which is characterized by a closed vulva and absence of oocytes (Figure S2D-E).

Next, to precisely evaluate the role of CSR-1, we measured small RNA accumulation (sRNA-seq), transcription (GRO-seq), mRNA stability (RNA-seq), and translation (Ribo-seq) in WT and mutant worms. In addition, to assess the direct effect of CSR-1 22G-RNAs on these processes, we sequenced the small RNAs bound to immunoprecipitated CSR-1 from similarly sorted late L4 worms to precisely identify the CSR-1 targets at the same developmental stage. We detected a total of 4803 genes with antisense 22G-RNAs loaded into CSR-1 (IP over input ≥ 2-fold enrichment and RPM ≥ 1 in each replicate of CSR-1 IP) (Table S1). The CSR-1 catalytic mutant displayed a global loss of 22G-RNAs for the majority of CSR-1 targets (Figure 1A, C). However, only 7.7% (n=119) of CSR-1 targets with >2-fold reduction of 22G-RNAs (n=1536) showed increased mRNA levels (Figure 1B), indicating that most mRNA targets are not destabilized by CSR slicer activity. The increase in mRNA and translational levels of the targets correlated with 22G-RNA levels in CSR-1 IPs in a dose-dependent manner (Figure 1D, E) in agreement with a previous report (Gerson-Gurwitz et al., 2016), but their transcription was unaffected (Figure 1F). Therefore, we conclude that CSR-1 slices a subset of target mRNAs having abundant 22G-RNAs. Moreover, we found that the targets that are post-transcriptionally regulated by CSR-1 are enriched for mRNAs encoding CSR-1 interacting proteins, identified by mass spectrometry (MS/MS) (enrichment factor 6.1, p < 1.6e^-28^) as direct, RNA-independent interactions (Figure S2G-I). Thus, CSR-1 slicer activity negatively regulates the expression of its own interactors, including CSR-1, suggesting a negative feedback loop.

**Figure 1.**
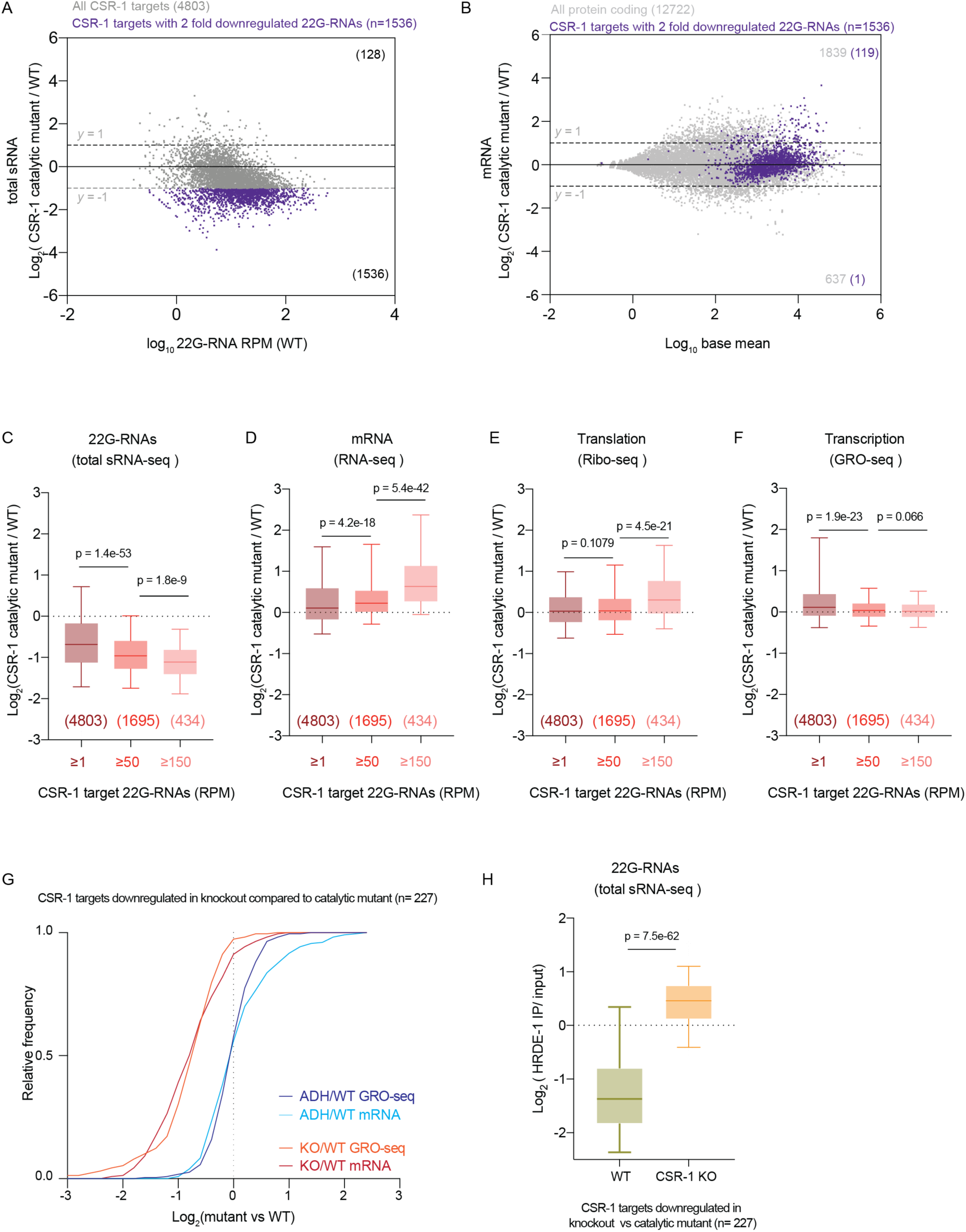
Defects in CSR-1 catalytic activity mainly impacts 22G-RNA abundance. **(A)** MA-plot showing total 22G-RNA log2 fold change between CSR-1 ADH (catalytic mutant) and WT. The number in parenthesis indicates the number of misregulated genes ≥ 2-fold. The average from two biologically independent replicates is shown. **(B)** MA-plot showing mRNA log2 fold change between CSR-1 ADH and WT. Genes with 22G-RNAs with 2-fold downregulation in CSR-1 catalytic mutant compared to WT are highlighted in purple. The average from two biologically independent replicates is shown, with “base mean” computed using DESeq2 (Love et al., 2014). **(C-F)** Box plots showing the log2 fold change of total 22G-RNAs (sRNA-seq), n = 2 biologically independent experiments **(C)**; or mRNAs (RNA-seq), n = 2 biologically independent experiments (**D**); mRNAs engaged in translation (Ribo-seq), n = 3 biologically independent experiments (**E**); and nascent RNAs (GRO-seq), n = 2 biologically independent experiments **(F)**, in CSR-1 ADH compared to WT strain. The distribution for the CSR-1 targets with 22G-RNA in CSR-1 IP ≥ 1 RPM, ≥ 50 RPM, or ≥ 150 RPM is shown (gene list in Table S1). Box plots display median (line), first and third quartiles (box), and 5^th^ /95^th^ percentile value (whiskers). **(G)** Cumulative frequency distribution for CSR-1 targets downregulated in CSR-1 KO compared to the CSR-1 ADH in GRO-seq (gene list in Table S1). The comparison shows GRO-seq (p = 1.6e^-49^) and RNA-seq (p = 4.2e^-37^) for CSR-1 KO or CSR-1 ADH compared to WT. **(H)** Box plots as in (C) showing the log2 fold change of 22G-RNAs (sRNA-seq) in HRDE-1 IPs compared to input in WT, CSR-1 KO (n = 2 biologically independent experiments). (Related figures-S2 and S3).

Overall, these results suggest that the main role of CSR-1 catalytic activity is to control the accumulation of 22G-RNAs. In addition, CSR-1 post-transcriptionally regulates a small fraction of CSR-1 targets that have highly abundant 22G-RNAs.

### CSR-1 protects a subset of oogenic enriched targets from piRNA-mediated transcriptional silencing

Similar to CSR-1 ADH worms, CSR-1 KO worms displayed a loss of 22G-RNAs as well as an upregulation of a subset of target mRNAs characterized by high abundance of 22G-RNAs (Figure S3A-D). However, the level of upregulation of CSR-1 target mRNAs was significantly lower in the *csr-1* KO compared to the *csr-1* ADH, possibly due to decreased transcription (Figure S3E). Indeed, we found that a subset of target genes displayed downregulated transcription and reduced mRNA levels in the KO compared to WT, but these were unaffected in the CSR-1 ADH (Figure 1G). The majority of these genes (53 %) were enriched for oogenic mRNAs (see Table S1 for gene list). Given that CSR-1 is proposed to protect germline transcripts from piRNA-mediated silencing, we hypothesized that in the *csr-1* KO, piRNAs can trigger the loading of 22G-RNAs into the nuclear Argonaute HRDE-1 resulting in the reduced transcription of this subset of CSR-1 targets. Indeed, HRDE-1 loads 22G-RNAs from transcriptionally downregulated CSR-1 targets in the *csr-1* KO (Figure 1H).

Overall, these data suggest a non-catalytic role of the CSR-1 protein in protecting a subset of oogenic targets from piRNA-mediated HRDE-1 transcriptional silencing.

### CSR-1 catalytic activity is required for biogenesis of 22G-RNAs on the coding sequence of target mRNAs

The global reduction of CSR-1-bound 22G-RNAs observed in CSR-1 mutants suggests that CSR-1 catalytic activity is required for 22G-RNA loading or biogenesis. Immunoprecipitation with WT or CSR-1 ADH proteins did not show any loss of binding of 22G-RNAs (Figure S4A), suggesting that catalytic inactive CSR-1 can still bind the 22G-RNAs produced in the mutant. We then investigated the distribution of CSR-1-bound 22G-RNAs along the target gene bodies. We found that 22G-RNAs are synthesized only at 3’ untranslated region (3’UTR) of their target mRNAs in the *csr-1* catalytic mutant, indicating that the RdRP fails to synthesize 22G-RNAs on the coding sequence (CDS) (Figure 2A-C). The CSR-1 catalytic mutation does not impair the loading of these 22G-RNAs generated from 3’UTR (Figure 2A, C and S4B). However, it fails to produce and load small RNAs from CDS (Figure 2A-C and S4B).

**Figure 2.**
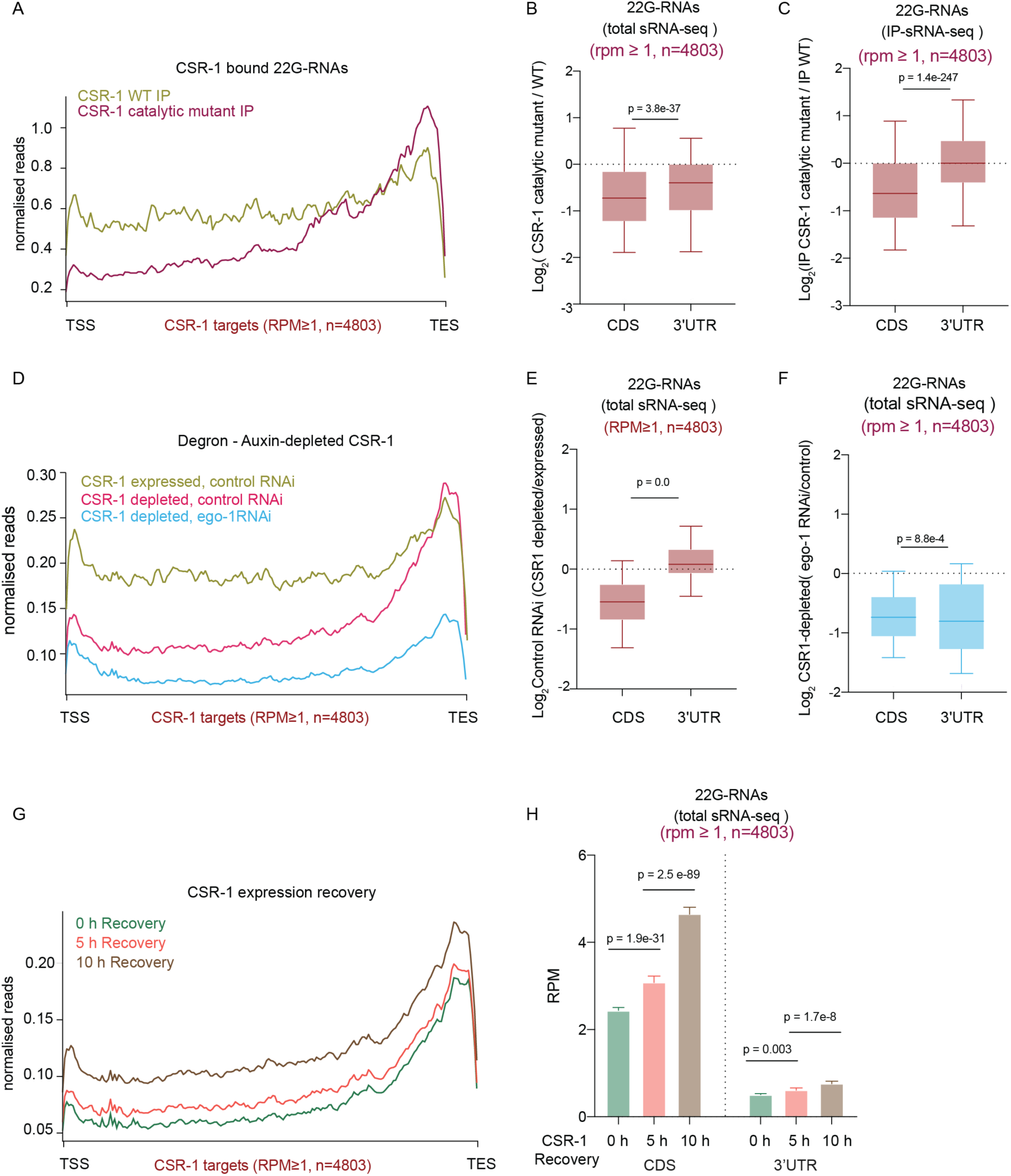
CSR-1 catalytic activity is required for the biogenesis of 22G-RNAs on the coding sequence. **(A)** Metaprofile analysis showing the distribution of normalized 22G-RNA (sRNA-seq) reads (RPM) across all CSR-1 targets (22G-RNA ≥1 RPM) in WT CSR-1 or CSR-1 ADH immunoprecipitation (IP). TSS indicates the transcriptional start site, TES indicates the transcriptional termination site. **(B)** Box plots showing the log2 fold change of the amount of total 22G-RNA generated from CDS and 3’ UTR of CSR-1 targets (22G-RNA ≥1 RPM) in CSR-1 ADH compared to WT. **(C)** Box plots showing the log2 fold change of the amount of 22G-RNA generated from CDS and 3’ UTR of CSR-1 targets (22G-RNA ≥1 RPM) bound in CSR-1 ADH IP compared to WT CSR-1 IP. Box plots display median (line), first and third quartiles (box), and 5^th^ /95^th^ percentile value (whiskers). Two-tailed P values were calculated using Mann–Whitney–Wilcoxon tests. n = 2 biologically independent experiments. **(D)** Metaprofile analysis as in (A) showing the distribution of normalized total 22G-RNA reads (RPM) across CSR-1 targets (22G-RNA ≥1 RPM) upon *ego-1* RNAi and Control RNAi treated in Auxin-depleted CSR-1 degron background (CSR-1 depleted) and degron control (CSR-1 expressed). **(E)** Box-plot as in (B) showing the log2 fold change in the amount of 22G-RNA generated from CDS and 3’ UTR of CSR-1 targets (22G-RNA ≥1 RPM) in Auxin-depleted CSR-1 compared to non-depleted CSR-1 degron control in control RNAi background. **(F)** Box-plot showing the log2 fold change in the amount of 22G-RNA generated from CDS and 3’ UTR of CSR-1 targets (22G-RNA ≥1 RPM) in *ego-1* RNAi compared to control RNAi treated in Auxin-depleted CSR-1 degron background, n = 2 biologically independent experiments. **(G)** Metaprofile analysis as in (A) showing the distribution of normalized total 22G-RNA reads (RPM) across CSR-1 targets (22G-RNA ≥1 RPM) after depletion of CSR-1, in the CSR-1 degron strain for 38 h by growing on auxin containing plates and recovery of CSR-1 expression by transferring on plates without auxin for 0 h, 5 h, and 10 h. **(H)** Bar graph representing the data in (F) showing the median RPM of 22G-RNAs with 95 % confidence interval generated from CDS and 3’ UTR of CSR-1 targets (22G-RNA ≥1 RPM) for CSR-1 expression recovered for 0, 5 or 10 h. Two-tailed P values were calculated using Mann–Whitney–Wilcoxon tests, n = 2 biologically independent experiments. (Related figure S4).

The RdRP EGO-1 has been proposed to exclusively synthesize CSR-1-bound 22G-RNAs (Claycomb et al., 2009; Gu et al., 2009; Maniar and Fire, 2011). To understand whether the small RNAs produced on the 3’UTR in the absence of CSR-1 protein or its catalytic activity are synthesized by EGO-1, we efficiently depleted CSR-1 using an auxin-induced degradation system, combined with *ego-1* RNAi knockdown (Figure S4C-E). First, we confirmed that CSR-1 22G-RNAs were depleted on CDS and enriched on 3’UTR upon auxin-induced CSR-1 depletion (Figure 2D, E), suggesting that the effect on CSR-1 22G-RNAs was not caused by a gain-of-function mutation in CSR-1 ADH. Next, we observed decreased 22G-RNAs from both CDS as well as 3’UTR upon *ego-1* RNAi knockdown (Figure 2D, F, S3F, G), implying that EGO-1 is exclusively responsible for the synthesis of the CSR-1 22G-RNAs in both WT and the *csr-1* mutants. However, the catalytic activity of CSR-1 is required to efficiently generate EGO-1-dependent 22G-RNAs along the coding sequences of target mRNAs.

Finally, we tested whether the restored expression of CSR-1 is sufficient to generate EGO-1-dependent 22G-RNAs on the gene body. For this purpose, we depleted CSR-1 by auxin-induced degradation for 38 h after hatching (0 h recovery) and then reintroduced CSR-1 by recovering expression for 5 and 10 hours (Figure S4H). As expected, the depletion of CSR-1 caused a loss of 22G-RNA accumulation on the CDS (Figure 2G and S4H - see 0 h recovery). However, upon reintroduction of CSR-1 expression (5 and 10 h recovery), we observed a steady increase of 22G-RNAs, mainly on the CDS (Figure 2G, H). The lack of complete recovery of 22G-RNAs could be due to the accumulation of germline defects as a result of CSR-1 depletion during the initial period of germline development.

Overall, these data demonstrate that EGO-1 can be recruited on the 3’UTR of target mRNAs and initiate the production of 22G-RNAs. However, CSR-1-mediated slicing of mRNAs is required to template the production of small RNAs on the gene body.

### Biogenesis of CSR-1 22G-RNAs and the regulation of their targets occurs in the cytosol

PIWI and RNAi biogenesis factors are known to localize in perinuclear condensates, called germ granules, and these germ granules have been proposed to be the site for biogenesis of 22G-RNAs (Chen et al., 2005; Kim et al., 2005; Phillips et al., 2012; Tops et al., 2005). CSR-1 and EGO-1 localize in both cytosol and the germ granules (Claycomb et al., 2009), suggesting that the biogenesis of CSR-1 22G-RNAs might also occur in these germ granules. To test this possibility, we used RNAi to simultaneously deplete four core components of germ granules (*pgl-1, pgl-3, glh-1*, and *glh-4*), called P granules (Figure S5A) (Updike et al., 2014). This treatment was sufficient to disrupt the localization of the GLH-1, PIWI and CSR-1 in germ granules (Figure 3A). However, the cytosolic localization of CSR-1 remained unaffected (Figure 3A).

**Figure 3.**
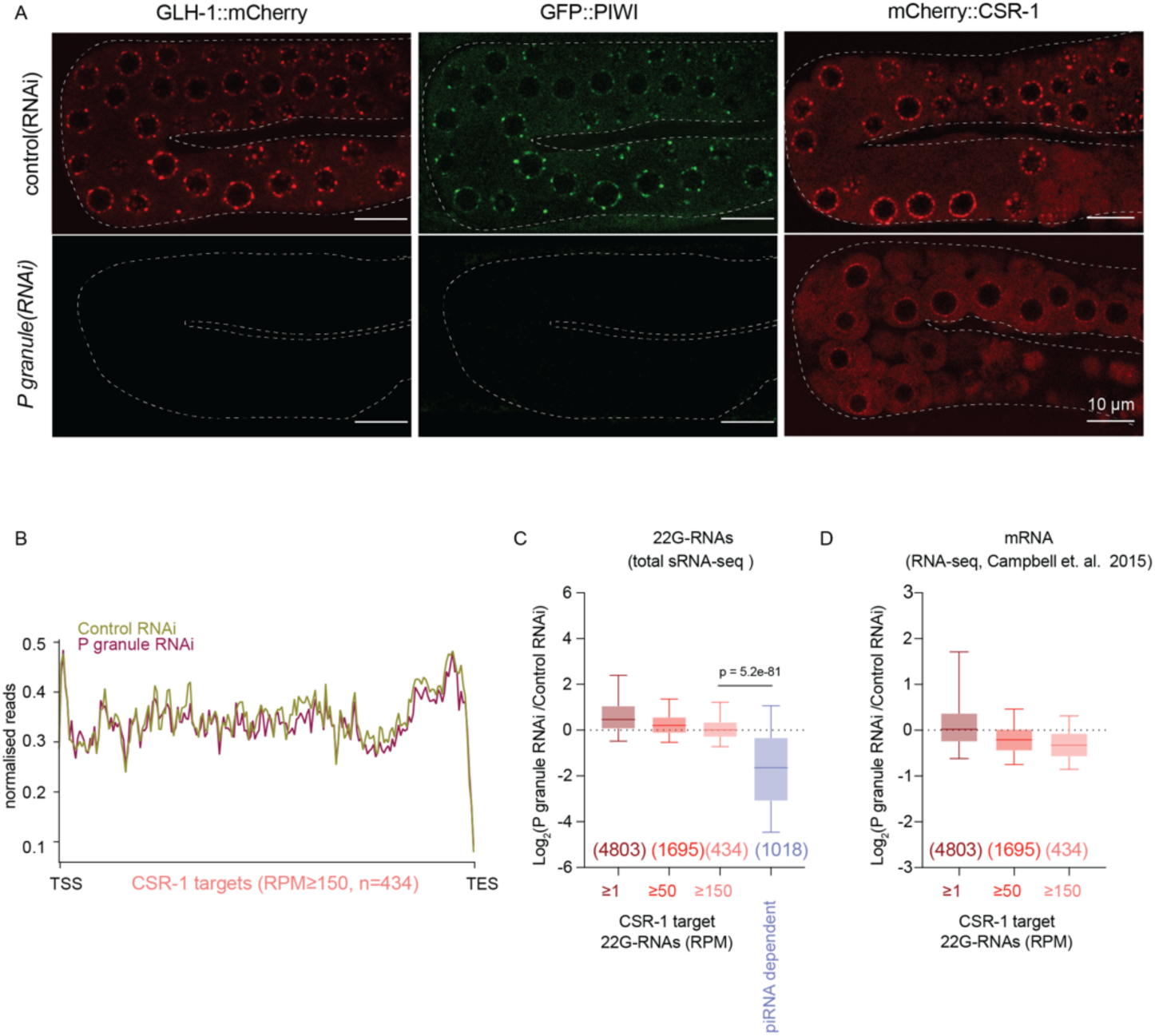
Biogenesis of CSR-1 22G-RNAs and the regulation of their targets do not require germ granule localization. **(A)** Fluorescent images of animals expressing P granule marker GLH-1:mCherry, GFP:PIWI or mCherry:CSR-1 treated with either control RNAi or P granule RNAi (*pgl-1, pgl-3, glh-1*, and *glh-4)*. CSR-1 is localized in both cytosol and P granule, and upon RNAi treatment, P granule localization of CSR-1 is lost while maintaining cytosolic localization. **(B)** Metaprofile analysis showing the distribution of normalized 22G-RNA (sRNA-seq) reads (RPM) across CSR-1 targets (22G-RNA ≥ 150 RPM) upon control RNAi and P granule RNAi. TSS indicates the transcriptional start site, TES indicates the transcriptional termination site. **(C)** Box plots showing the log2 fold change of total 22G-RNA (sRNA-seq) upon P granule RNAi compared to control RNAi. The distribution for the CSR-1 targets with 22G-RNA in CSR-1 IP ≥ 1 RPM, ≥ 50 RPM, or ≥ 150 RPM and piRNA dependent 22G-RNA target genes (Barucci et al. 2020) are shown. Box plots display median (line), first and third quartiles (box), and 5^th^ /95^th^ percentile value (whiskers). Two-tailed P values were calculated using Mann–Whitney–Wilcoxon tests, n = 2 biologically independent experiments. **(D)** Box plots showing the log2 fold change of mRNA expression (RNA-seq from (Campbell and Updike, 2015)) upon P granule RNAi compared to control RNAi. The distribution for the CSR-1 targets with 22G-RNA in CSR-1 IP ≥ 1 RPM, ≥ 50 RPM, or ≥ 150 RPM is shown. (Related figure S5).

Next, we evaluated the effects of loss of germ granule localization of PIWI and CSR-1 on 22G-RNA biogenesis. As expected, piRNA-dependent 22G-RNAs were globally depleted upon P granule RNAi treatment (Figure 3C). Surprisingly, CSR-1 22G-RNAs were unaffected upon P granule RNAi treatment, despite the loss of perinuclear CSR-1 germ granule localization (Figure 3B, C). Furthermore, CSR-1 targets were not upregulated upon P granule RNAi (RNA-seq data from (Campbell and Updike, 2015) (Figure 3D and S5A). These results highlight that CSR-1 22G-RNA biogenesis occurs in the cytosol independently of germ granules.

### Translating mRNAs serve as the template for 22G-RNA biogenesis

Our data so far suggest that CSR-1 22G-RNAs are generated in the cytosol. Consistent with CSR-1 localization in the cytosol and germ granules, we identified ribosomal and ribosomal-associated proteins, which are enriched in the cytosol, and germ granule components in our IP-MS/MS as direct CSR-1 interactors (Figure 4A and (Barucci et al., 2020). Moreover, CSR-1 ADH showed reduced co-purification of ribosomal proteins and increased co-purification of germ granule components, compared to CSR-1 WT (Figure 4B) and localizes primarily in germ granules (Figure S5B).

**Figure 4.**
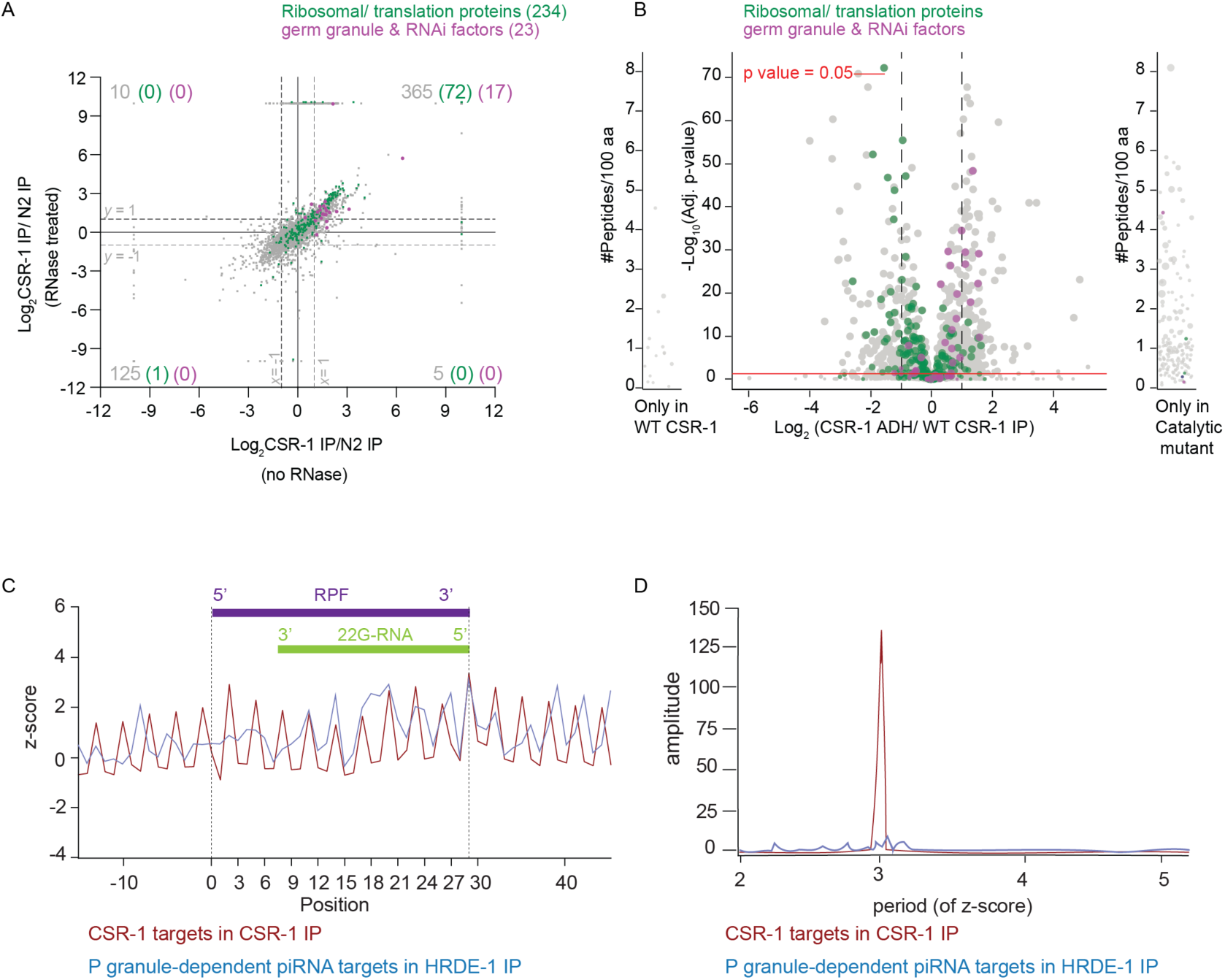
CSR-1 22G-RNAs are synthesized concomitantly with mRNA translation. **(A)** Scatter plot comparing the log2 fold changes in CSR-1 interactors (IP-MS/MS) to control IPs performed in WT strain in the absence of RNase treatment (x-axis) to the IPs performed after RNase treatment ((Barucci et al., 2020) and this study) (Table S2). Ribosomal proteins and translation regulators are highlighted in green, and germ granule proteins, including RNAi factors, are highlighted in magenta. Number in grey refers to all interactors with log2 fold change of ≥1 and p-value ≤ 0.05 for each quadrant. The number in parenthesis is for ribosomal and translation associated proteins enriched, and granule & RNAi factors. n = 4 biologically independent experiments. **(B)** Volcano plot showing log2 fold change in enrichment values and corresponding significance levels for proteins co-purifying with CSR-1 ADH compared to WT CSR-1 (Table S3). Ribosomal proteins and translation regulators are highlighted in green. Germ granule proteins, including RNAi factors, are highlighted in magenta. The size of the dots is proportional to the number of peptides used for the quantification. The linear model was used to compute the protein quantification ratio, and the red horizontal line indicates the two-tailed p-value = 0.05. n = 4 biologically independent experiments. **(C)** Plot showing the z-score for read density for the of 5’ terminus of 22G-RNAs for CSR-1 targets (RPM≥1) in CSR-1 IP and P granule dependent piRNA targets in HRDE-1 IP (gene list in Table S1) relative to the start of 29-nt long Ribosomal protected fragments (RPF). **(D)** Periodogram based on Fourier transform for read-density around RPF 5’ start position showing periodicity of CSR-1 22G-RNAs phasing with RPFs. P granule dependent piRNA targets in HRDE-1 IP was used as control (Related figure S5).

Based on these data, we hypothesized that 22G-RNAs are synthesized in the cytosol, using translating mRNAs as templates. To test this hypothesis, we mapped the distance between the start of the 29-nucleotide Ribosomal Protected Fragments (RPF) (Aeschimann et al., 2015) and the 5’ end of CSR-1 22G-RNAs (Figure 4C). We observed periodicity of 3 nucleotides in phase with ribosomes (Figure 4D), indicating that the synthesis of CSR-1 22G-RNAs occurs on mRNA templates engaged in translation. In contrast, the HRDE-1 loaded 22G-RNAs of P granule dependent piRNA targets (Table S1) did not show phasing with ribosomes as observed due to a lack of 3 nucleotide periodicity (Figure 4C and D), in agreement with the fact that P granules are devoid of translating mRNAs (Lee et al., 2020; Schisa et al., 2001).

Altogether these results suggest that CSR-1 cleaves actively translating mRNAs, which become the template for EGO-1-mediated synthesis of 22G-RNAs on the coding sequence of mRNA targets.

### mRNA translation antagonizes CSR-1 22G-RNA biogenesis

EGO-1 mediated synthesis of CSR-1 22G-RNAs does not occur on every germline mRNA at similar levels, and we found that the levels of 22G-RNA are independent of the levels of the mRNA template (Figure S6A). Given our observations that actively translating mRNAs serve as the template for CSR-1 22G-RNAs, we hypothesized that the translation efficiency (TE) of germline mRNAs impacts CSR-1 22G-RNA biogenesis. To test this hypothesis, we calculated the TE of CSR-1 targets using the Ribo-seq and RNA-seq data from WT worms at the late L4 stage. We observed that levels of CSR-1 associated 22G-RNAs produced from a target mRNA were inversely correlated with their TE (Figure 5A), suggesting that translation antagonize the biogenesis of CSR-1 22G-RNAs.

**Figure 5.**
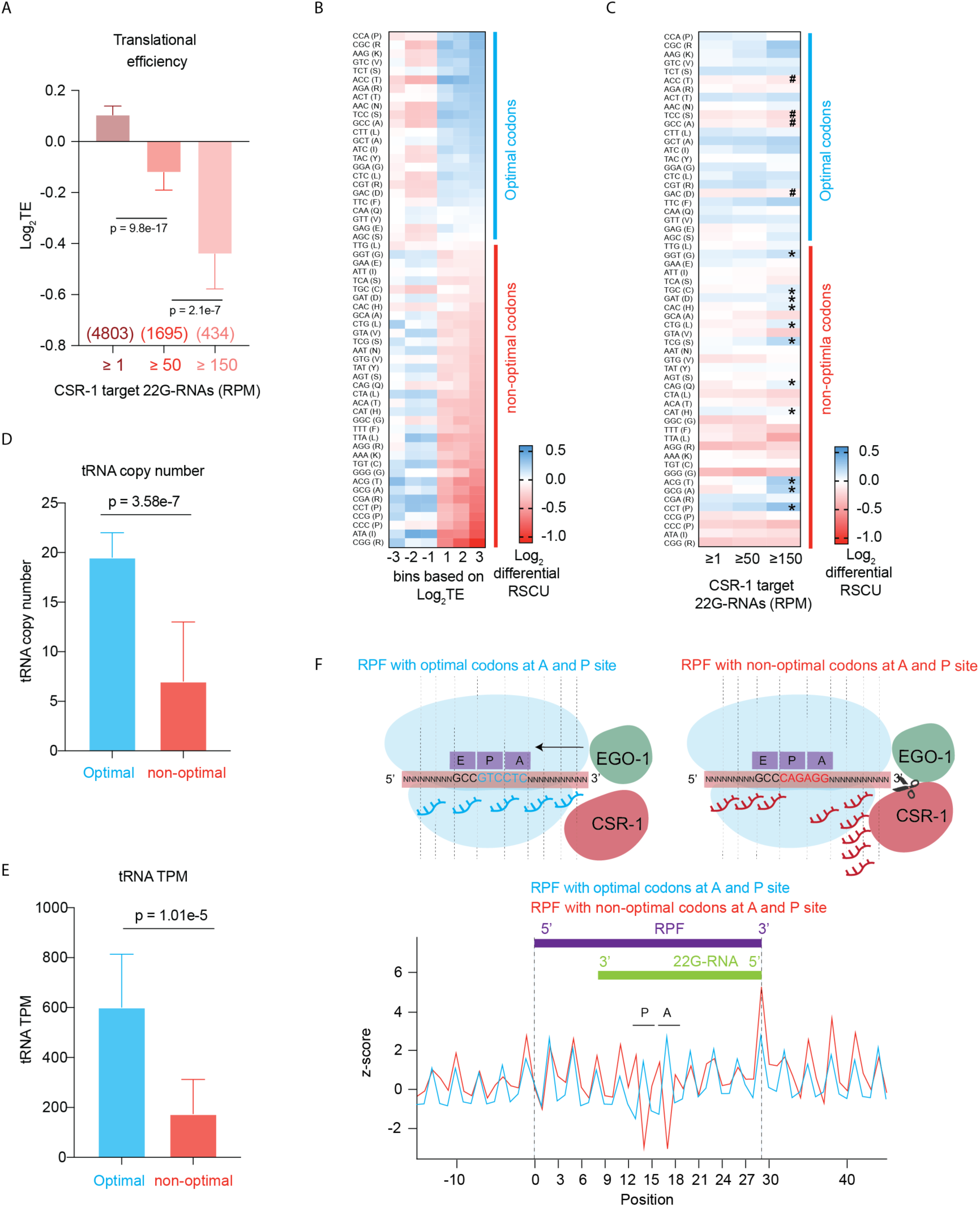
Highly translated mRNAs and optimal codons negatively correlate with CSR-1 22G-RNA abundance. **(A)** Translation efficiency Log2 (RPF TPM/ mRNA TPM) for CSR-1 targets in WT strain. The distribution for the CSR-1 targets 22G-RNA in CSR-1 IP ≥ 1 RPM, ≥ 50 RPM, or ≥ 150 RPM is shown. Bars represent median with 95 % confidence interval. Two-tailed P values were calculated using Mann– Whitney–Wilcoxon tests, n = 2 biologically independent experiments. **(B)** heat map showing log2 fold change in Relative Synonymous Codon Usage (RSCU) for all protein-coding genes categorized by increasing translational efficiency compared to genes showing neutral translational efficiency of 1, as explained in methods. Codons are arranged in order of decreasing RSCU (top to bottom) in the category of log2 TE ≥3. The blue line highlights optimal codons in genes with high TE, and the red line highlights non-optimal codons. **(C)** heatmap similar to (B) showing log2 fold change in Relative synonymous codon usage for all CSR-1 targets (≥ 1, ≥ 50, and ≥150 RPM of 22G-RNA) compared to genes showing neutral translational efficiency as explained in methods. **‘**\*’ marks over-used non-optimal codons by CSR-1 targets and ‘#’ marks under-used optimal codons. **(D)** Bar graph showing the median copy numbers for tRNAs for optimal or non-optimal codons with a 95 % confidence interval, **(E)** Bar graph showing the median TPMs for tRNAs from the GRO-seq dataset for WT strain at the late l4 larval stage (44h) for optimal or non-optimal codons with a 95 % confidence interval. For codons with missing cognate tRNA, values have been adjusted by considering tRNA copy numbers and TPMs for tRNA recognizing these codons by wobble base pairing (Reis et al., 2004). (see Figure S5B, C for non-adjusted values). Two-tailed P values were calculated using Mann–Whitney–Wilcoxon tests, n = 2 biologically independent experiments. **(F)** Plot showing the z-score for the read density for the of 5’ terminus of 22G-RNAs for CSR-1 targets (RPM≥150) relative to the start of 29-nt long Ribosomal protected fragments (RPF) with either optimal or non-optimal codons at their P and A site. The scheme shows possible initiation of 22G-RNA biogenesis after a slicing event downstream of RPF with bad codons at the A and P site. (Related figure S6).

Codon usage and the availability of the tRNA pool influence TE (Pechmann and Frydman, 2012; Presnyak et al., 2015). Therefore, we investigated whether these mechanisms affect the biogenesis of CSR-1 22G-RNAs. We determined optimal and non-optimal codons using our experimental data from Late L4 staged worms. First, we calculated the normalized average relative synonymous codon usage (RSCU) for genes for different categories of high or low TE (Figure 5B). Codons showing enrichment in genes with high TE (log2TE ≥3) were considered optimal codons, and the ones under-represented were considered non-optimal codons (Figure 5B). We confirmed that our classification of optimal/non-optimal codons correlated with tRNA copy number (Figure 5D, S6B, D) and tRNA pool available in the late L4 worm population (44h) as measured by GRO-seq (Figure 5E, S6C, E). We noticed that for codons with no tRNA cognates and requiring tRNA binding by wobble pairing, all optimal codons end with C, and non-optimal with U. Translation elongation is lower for those ending with a U (Stadler and Fire, 2011).

We then evaluated the codon usage of CSR-1 targets by comparing their normalized average RSCU to highly translated mRNAs. We found that non-optimal codons were enriched and optimal codons were depleted in CSR-1 targets, suggesting that this might be an encoded feature of mRNA targets influencing the priming of 22G-RNA synthesis (Figure 5C). Non-optimal codons are known to promote ribosome stalling (Duret, 2000; Novoa and Pouplana, 2012; Tuller et al., 2010). To map differences in 22G-RNA biogenesis on sequences with optimal or non-optimal codons, we divided RPFs into two categories based on the presence of either an optimal or non-optimal codon at the A and P sites of the ribosome and then mapped the distance between 5’ of 22G-RNAs and RPFs. We observed a peak for the 5’ end of 22G-RNAs downstream of RPF (29^th^ position) when the A and P sites of the ribosomes are occupied by a non-optimal codon contrary to when optimal codons are present on A and P sites which show no bias (Figure 5F). This result suggests that the 22G-RNA production is preferentially initiated downstream of ribosomes especially occupying bad codons, by CSR-1 mediated slicing and recruitment of EGO-1.

Altogether, these observations suggest that translation and ribosome position dictate the production of CSR-1 22G-RNAs.

### Increasing the translation efficiency of CSR-1 target impairs CSR-1 22G-RNA biogenesis and function

To determine whether non-optimal codons directly affect TE and CSR-1 22G-RNA biogenesis, we altered the coding potential of a CSR-1 target. We examined *klp-7*, which has the second-highest abundance of 22G-RNAs loaded by CSR-1 and is post-transcriptionally regulated by CSR-1.

KLP-7 is a kinesin-13 microtubule depolymerase and is required for spindle organization and chromosome segregation (Gigant et al., 2017). Overexpression of KLP-7 in the *csr-1* mutant has been shown to cause microtubule assembly defects (Gerson-Gurwitz et al., 2016). *klp-7* showed enrichment of non-optimal codons and depletion of optimal codons similarly to other CSR-1 targets (Figure S7A). We optimized the codon usage in *klp-7* by incorporating exclusively synonymous optimal codons (Figure S7A). We used CRISPR-Cas9 to replace endogenous *klp-7* isoform b with the modified *klp-7* codon-optimized (*klp-7_*co) to avoid disrupting potential UTR-mediated regulation.

To ascertain whether codon optimization of *klp-7* affected the TE, we performed RNA-seq and Ribo-seq from synchronized and sorted late L4 population (44 h). Indeed, we detected a 2-fold increase in the TE of *klp-7* mRNA in the *klp-7*_co strain compared to WT (Figure 6A). The TE of other CSR-1 targets remained unaffected in *klp-7*_co strain, indicating that the effects observed are specific to *klp-7* mRNA (Figure 6A). In addition, KLP-7 protein levels were increased in two independent lines of *klp-7*_co compared to WT, consistent with increased translation (Figure S7C). We then measured the level of 22G-RNAs antisense to *klp-7* mRNA in the *klp-7*_co strain compared to WT, and we observed a 1.4-fold decrease in 22G-RNAs (Figure 6A). The levels of 22G-RNAs for other CSR-1 targets remained unaffected (Figure 6A). Further, the significant decrease in 22G-RNAs on the optimized *klp-7*_co allele was observed in exons 3-6 and was accompanied by an increase in ribo-seq peak height at those positions (Figure 6B). The *klp-7*_co strain also showed increased *klp-7* mRNA level compared to WT (Figure 6A), and we confirmed this result by RT-qPCR (Figure 7B). These results suggest that CSR-1 targeting and regulation is impaired on *klp-7_co* mRNA. To validate this, we performed *csr-1* RNAi and showed increased *klp-7* Mrna levels in the WT strain but not in the *klp-7*_co strain (Figure 6C), suggesting that CSR-1 activity is reduced at *klp-7*_co mRNA. The increased levels of *klp-7* mRNA correlated with a reduction in brood size (Figure S7D) and higher embryonic lethality at 25°C in *klp-7*_co strain compared to WT (Figure S7E), indicating the physiological relevance of *klp-7* mRNA targeting by CSR-1. Finally, to rule out any difference in the production of either 22G-RNAs or mRNA levels due to possible developmental defects between *klp-7*_co and WT strain, we generated a heterozygote strain of *klp-7*_co with a fluorescent GFP marker on the balancer chromosome. We sorted heterozygote GFP positive worms with one copy of modified *klp-7_co* and one copy of WT *klp-7* each and performed RNA-seq and sRNA-seq. We observed similar results with a 1.8-fold increase in mRNA levels for *klp-7*_co compared to the WT *klp-7* copy and a 1.25-fold decrease in 22G-RNA levels (Figure S6F-G). These results demonstrate that CSR-1 22G-RNA biogenesis and activity is reduced on mRNA templates with optimized codons and increased translation.

**Figure 6.**
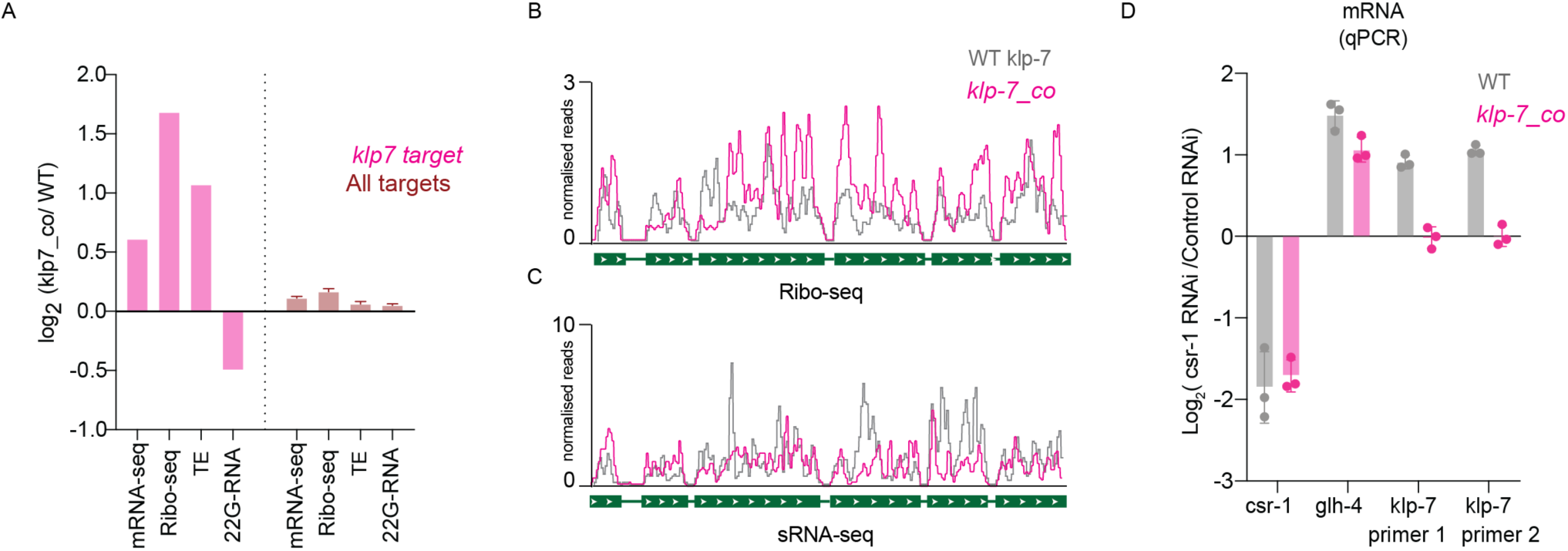
Increasing optimal codon usage of CSR-1 target decreases small RNA production. **(A)** Plot showing log2 fold change for normalized reads for mRNAs from RNA-seq, RPF from Ribo-seq, and 22G-RNAs from sRNA-seq and differential Translational efficiency for *klp-7* (top CSR-1 target) and all CSR-1 targets (RPM≥1, n=4803) for the strain with codon-optimized *klp-7* (*klp7_co*) compared to WT strain. n = 2 biologically independent experiments. **(B, C)** A genomic view of *klp-7* showing of Ribo-seq **(B)** and 22G-RNAs **(C)**, normalized reads for the strain with codon-optimized *klp-7* (*klp7_co*) compared to WT strain. **(D)** log2 fold change in expression of *csr-1, glh-4*, and *klp-7* upon *csr-1* RNAi compared to control RNAi by qPCR in the WT strain and *klp-7_co* (strain with *klp-7* codon optimization). n = 3 biologically independent experiments. (Related figure S7).

**Figure 7.**
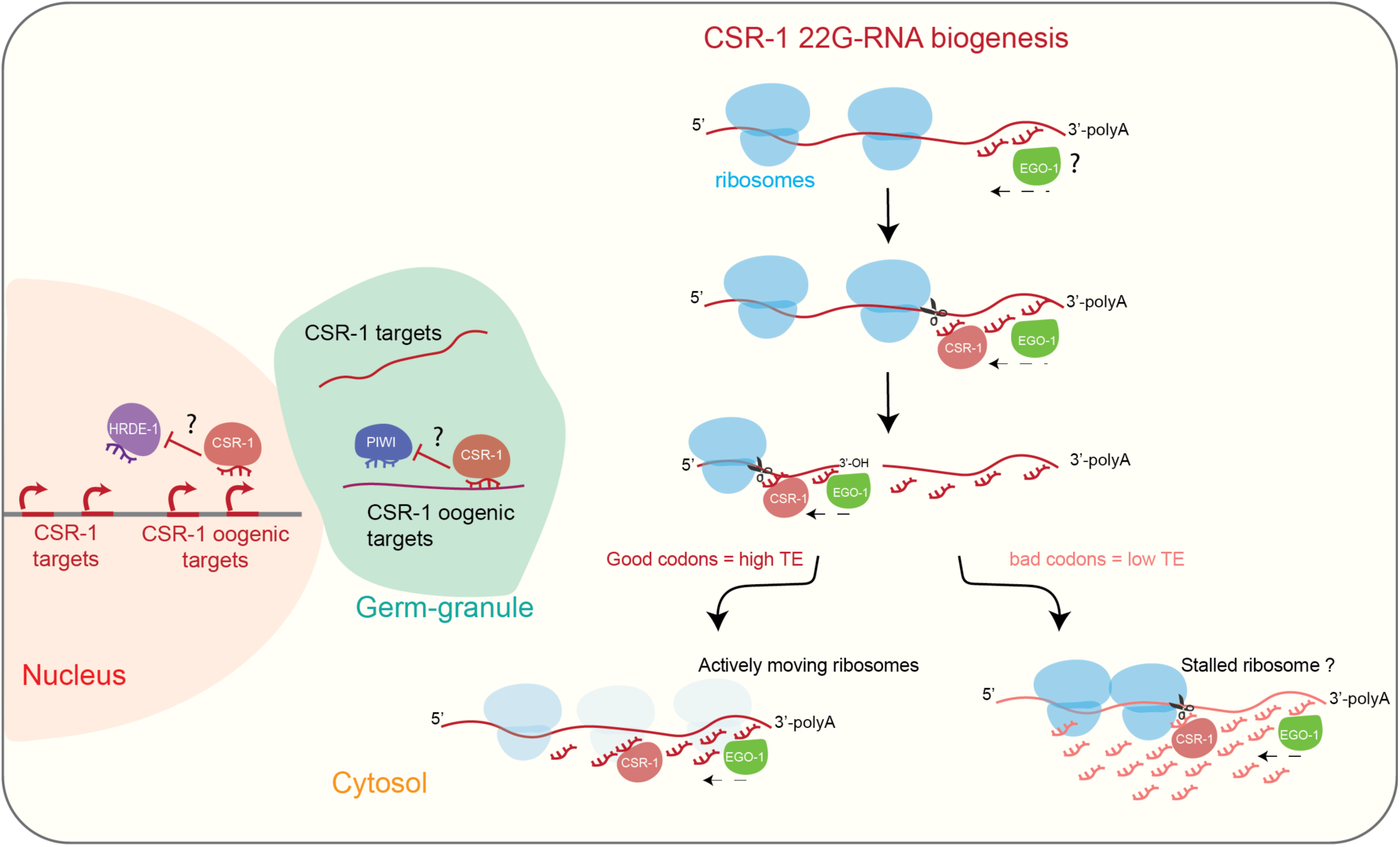
Model illustrating biogenesis of CSR-1-22G-RNA in the cytosol. CSR-1 targets most of the germline expressed genes. CSR-1 22G-RNAs are produced from mRNAs engaged in translation in the cytosol. We propose EGO-1 initiate 22G-RNA biogenesis at the 3’UTR on every actively translating mRNAs or by being recruited on specific 3’UTR sequences by yet unknown mechanism. However, to produce 22G-RNAs on coding sequence, CSR-1 slicing activity is required on the mRNA template. Codon usage and translation efficiency antagonistically regulate levels of 22G-RNAs production on different CSR-1 targets. We propose that CSR-1 can interact with ribosomes and the slicing event is more biased downstream of a possible stalled ribosome occupying a bad codon site. CSR-1 slicer activity can regulate gene-expression of few top targets, which further depends on the 22G-RNA levels. Additionally, CSR-1 can protect a set of mainly oogenic genes from piRNA mediated HRDE-1 transcriptional silencing, in a catalytic independent manner

Altogether, these results suggest efficiently translating ribosomes block the access of CSR-1 to the mRNA template and thereby hamper 22G-RNA production and, in turn, affects gene regulation by CSR-1 during germline development.

## DISCUSSION

In this study, we have determined the rules governing germline mRNA targeting by CSR-1 and addressed the long-standing paradox of CSR-1 function as anti-silencer or a slicer. We show a significant fraction of the slicing activity of CSR-1 is directed towards the production of 22G-RNAs on the coding sequence of its targets. We demonstrate that only a fraction of CSR-1 slicing activity regulates targets post-transcriptionally in the germline.

Our observations also suggest a catalytic-independent function of CSR-1 in preventing piRNA-dependent chromatin silencing. Specifically, we showed that in the absence of CSR-1 protein, a subset of CSR-1 target genes is misrouted into the piRNA pathway, which represses their expression at transcriptional levels through the nuclear Argonaute HRDE-1. Therefore, in addition to the post-transcriptional regulation of germline mRNAs (Gerson-Gurwitz et al., 2016), CSR-1 can also license the transcription of germline genes.

We further dissected the mechanism of CSR-1 22G-RNA production. We demonstrated that the synthesis of 22G-RNAs occurs in the cytosol on translating mRNA templates, whereas germ granules are dispensable for CSR-1 regulation. We propose that a low translation efficiency favors the CSR-1 slicer activity on the target mRNA occupied by ribosomes and initiates 22G-RNA biogenesis by priming RdRP EGO-1 activity. Finally, we have determined how CSR-1 can preferentially target some germline mRNAs. We discovered that incorporation or avoidance of non-optimal codons is a strategy adopted by germline mRNAs to be differentially regulated by CSR-1 22G-RNAs.

Overall, this study highlights the codon dependence and translational efficiency of mRNAs in the germline for the regulation of CSR-1-22G-RNAs biogenesis and, in turn, gene expression of the targets, which could have a significant bearing on germline gene regulation not just in worms but across species.

### Biogenesis of CSR-1 22G-RNAs

In this study, we have established that the Argonaute CSR-1 slices target mRNAs to trigger the generation of RdRP-dependent 22G-RNAs on the gene body. We propose that CSR-1 slicer activity is required to generate new 3’-OH ends along the gene transcript to facilitate the initiation of RdRP (EGO-1) synthesis towards the 5’ end of the mRNA target. This is consistent with previous *in vitro* RdRP analysis showing that non-polyadenylated 3’-OH ends of RNAs served as better substrates for 22G-RNA synthesis (Aoki et al., 2007), suggesting that the cleavage of RNA may be vital for the processivity of RdRPs. Based on these results, we speculate that no primary small RNAs are required to generate CSR-1 22G-RNAs along the mRNA sequence. Instead, CSR-1 catalytic activity triggers the synthesis of 22G-RNAs by the RdRP EGO-1, starting from the 3’UTR of the target transcripts. Even if the catalytic activity of CSR-1 is required to generate 22G-RNAs along the gene body of target transcripts, it is still unknown what triggers the recruitment of EGO-1 on the 3’UTR. Primary small RNAs, which are yet to be identified, might prime the activity of EGO-1. Alternatively, EGO-1 might produce low levels of 22G-RNAs from the polyadenylated tail of mRNAs instead of cleaved 3’OH end products. Thus, these low levels of 22G-RNAs, which are then loaded into CSR-1 can initiate the production of 22G-RNAs along the gene body. RNA binding proteins and/or other unknown factors together with specific sequences in the 3’UTR might also recruit and initiate EGO-1-dependent 22G-RNAs from the 3’UTR of selected mRNAs.

### The role of translation and codon usage in CSR-1 22G-RNA biogenesis

Most germline-expressed mRNAs are targeted by CSR-1 to promote EGO-1 dependent 22G-RNAs, but they generate different levels of 22G-RNAs. We found that CSR-1 targets in adult germlines are mainly actively translating mRNAs, and 22G-RNAs are synthesized in phase with ribosomes. Therefore, the CSR-1-22G-RNA biogenesis machinery likely needs to cope with the presence of ribosomes on the target transcripts. We show that non-optimal codons in germline mRNAs enhance the capacity of CSR-1 to prime the synthesis of EGO-1-dependent 22G-RNAs along the gene body. The use of non-optimal codons by CSR-1 targets and priming of 22G-RNAs at stalled positions is a way to cope with the ribosomal presence on the target transcripts. Therefore, sequences that promote ribosome stalling promote targeting by CSR-1. We demonstrate that substituting non-optimal codons with optimal codons is sufficient to allow germline mRNAs to escape CSR-1-dependent regulation. There is increasing evidence that the translation machinery associates with the Argonautes and small RNA biogenesis factors. Recently it was shown that RNAi can occur co-translationally with an accumulation of ribosomes upstream of the dsRNA targeted region (Pule et al., 2019). Another study proposed that ribosomes coordinate with piRNA biogenesis factors in mouse testes to achieve endonucleolytic cleavage of non-repetitive long RNAs to produce pachytene piRNAs (Sun et al., 2020). There is a further report suggesting possible repression of translation by 22-nt siRNAs in plants and induction of transitive small RNA amplification by RNA-dependent RNA polymerase 6 (RDR6) (Wu et al., 2020). Therefore, we propose that the regulation of small RNA biogenesis by ribosome occupancy and codon usage of the target transcript might be a general strategy adopted across evolution.

### Granule vs Cytosolic functions of CSR-1

We found that the slicer activity of CSR-1 and 22G-RNA biogenesis at germline mRNA targets are independent of germ granules. This raises the question on the function of CSR-1 in germline granules. CSR-1 might be enriched in germ granules of adult gonads to prevent CSR-1 slicer activity on the majority of germline mRNAs. Indeed, only 7.7% of CSR-1-dependent 22G-RNA targets are significantly regulated by CSR-1 slicer activity in adults. Moreover, the majority of these targets are CSR-1-interacting proteins suggesting a negative feedback regulation of the CSR-1 pathway. This is in contrast with the recently described function of the maternally delivered CSR-1 in the embryo, which exclusively localizes in the cytosol of somatic blastomere where it cleaves and clears hundreds of maternal mRNA targets (Quarato et al., 2020). Therefore, we propose that CSR-1 slicer activity on mRNA targets is partially suppressed in the germline by titrating away a part of CSR-1 in germ granules and primarily serves to generate interacting small RNAs in the cytosol that fully operates intra-generationally in the embryo. This also explains why despite targeting almost all germline genes, CSR-1 catalytic activity regulates the expression of only a few in the germline. In addition, CSR-1 localization in the germ granule might serve to antagonize piRNA-dependent targeting on germline mRNAs and therefore license those transcripts to be translated in the cytosol. Indeed, we have shown that most of the piRNA-dependent 22G-RNAs are generated in germ granules, and we propose that the competition between CSR-1 and PIWI might occur in germ granules.

## ACKNOWLEDGEMENTS

We would like to thank all the members of the Cecere laboratory, Manish Grover, Sudarshan Gadadhar, and Angela Anderson (Life Science Editors) for the helpful discussions on the paper. We thank Micheline Fromont for her help to set up Ribosome profiling. We thank Celine Didier for technical assistance. We thank the Heng-Chi Lee lab, Miska lab, Desai lab, Updike lab and Kennedy lab for sharing strains and reagents. Some strains were provided by the CGC, funded by the NIH Office of Research Infrastructure Programs (P40 OD010440).

This project has received funding from the Institut Pasteur, the CNRS, and the European Research Council (ERC) under the European Union’s Horizon 2020 research and innovation programme under grant agreement No ERC-StG-679243. MS and EC were supported by the

Pasteur-Roux-Cantarini Postdoctoral Fellowship program. PQ was supported by Ligue Nationale Contre le Cancer (SFB19032). FD and DL have received funding from Région Ile-de-France and Fondation pour la Recherche Médicale grants to support this study.

## AUTHOR CONTRIBUTIONS

GC and MS identified and developed the core questions addressed in the project. MS performed most of the experiments and analyzed the results together with GC. EC and LB generated all the lines used in this study with the help of MS and PQ. EC performed the imaging experiments, RNA-seq for CSR-1 mutants and IP-sRNA-seq of HRDE-1 in CSR-1 KO. PQ performed GRO-seq for CSR-1 mutants. BL performed all the bioinformatics analysis. SP contributed for distance mapping analysis of 22G-RNA reads and Ribo-seq reads with BL. FD and DL performed MS/MS experiments and analyzed the data together with MS. MS and GC wrote the paper with the contribution of EC and PQ.

## MATERIALS AND METHODS

### *C. elegans* strains and maintenance

Strains were grown at 20 °C on NGM plates seeded with *E. coli* OP50 using standard methods (Brenner, 1974) unless otherwise stated. The wild-type reference strain used was Bristol N2. A complete list of strains used in this study is provided in Table S4

### Generation of CRISPR–Cas9 lines

Cas9-guide RNA (gRNA) ribonucleoprotein complexes were microinjected into the hermaphrodite syncytial gonad as described previously (Paix et al., 2015) and gRNA design and *in vitro* synthesis were done following the protocol detailed in (Barucci et al., 2020). For the introduction of a *csr-1(D769A)* mutation (Barucci et al., 2020) in *3×flag::ha::csr-1* animals, we used a single-stranded oligonucleotide repair template ordered from IDT as standard 4 nM ultramer oligo. For the endogenous *klp-7* gene replacement, we used two gRNAs, each one respectively targeting a region at the 5’ and 3’ of the *klp-7* isoform b gene. A PCR repair template containing 33bp homology arms, was directly amplified from a plasmid containing a codon-optimized version of *klp-7* (*klp-7_co* synthetic gene) synthesized from GenScript (Table S5).

Mix concentrations were adapted from (Dokshin et al., 2018). In brief, 10 µl mixes typically contained the following final concentrations: 0,1µg/µL Cas9-NLS protein (TrueCut V2, Invitrogen), 100 ng/µl *in vitro* transcribed target-gene gRNA, 80 ng/µl of target-gene ssODN repair template or 300 ng/µl target-gene double-stranded DNA repair template and 80 ng/uL pRF4 (roller marker). Cas9 and the target-gene gRNA were pre-incubated 10-15 minutes at 37°C before the addition of the other components to the mixture. dsDNA repair templates were subjected to a melting/annealing step (Dokshin et al., 2018) before addition to the final mix. A detailed list of gRNAs, single-stranded DNA and double-stranded DNA repair templates and primers used for genotyping is provided in Table S5.

### RNAi

RNAi clone for *ego-1* and *csr-1* used in this study were obtained from the Ahringer library (Kamath and Ahringer, 2003). For quadruple P granule RNAi (*pgl-1, pgl-3, glh-1* and *glh-4)* pDU49 clone (gift from Updike lab (Updike et al., 2014)) was used. An empty vector (L4440) was used as a control in all of our RNAi experiments. RNAi experiments were performed by growing synchronous population of L1 larvae on Petri dishes with NGM and IPTG (15 cm) seeded with concentrated RNAi food. For *csr-1* and *ego-1* RNAi worms were grown from L1 to late L4 stage on RNAi food at 20°C. For P-granule RNAi worms were grown for two generations at 25°C (Updike et al., 2014). Post-RNAi treatment, worms were harvested and sorted on COPAS biosorter to enrich late L4 larvae. RNAi efficacy was confirmed by RT-qPCR.

### *ego-1* RNAi and auxin-induced CSR-1 degradation

For *ego-1* RNAi, worms were grown from L1 to 38 hours post-hatching on RNAi or control food on IPTG containing plates and then washed twice with M9 buffer and then shifted to either Auxin plates or Ethanol plates (containing 500 µM auxin, 0.5 % Ethanol or only 0.5 % Ethanol respectively) to deplete degron tagged CSR-1 by auxin-induced degradation as described before (Quarato et al., 2020). Plates were seeded with respective *ego-1* RNAi or control RNAi food. Auxin-induced degradation was performed for 6 h. Worms were then harvested, washed with M9 buffer and sorted on COPAS biosorter to enrich for Late L4 larval population. CSR-1 depletion was confirmed by live imaging.

### CSR-1 expression recovery post auxin-induced degradation

A synchronous population of degron-tagged CSR-1 strain was grown on NGM plates containing 500 µM auxin, 0.5 % ethanol from L1 to 38 hours post-hatching to degrade degron-tagged CSR-1. After 38 hours, worms were washed thrice with M9 buffer and divided into three parts. 1/3^rd^ worms were immediately sorted on COPAS biosorter to enrich for a synchronous population for 0 h recovery time point of CSR-1 expression. The rest of the worms were seeded on two NGM plates and allowed to grow in the absence of auxin induction for 5 or 10 h to recover CSR-1 expression. Worms were washed with M9 buffer at respective time points and sorted using COPAS biosorter to enrich for a synchronised population for each time point. CSR-1 expression was monitored using live imaging.

### Brood-size assay

For the brood size single L1 larvae were manually picked and placed onto NGM plates seeded with *E. coli* OP50 and grown at 20°C or 25°C until adulthood and then transferred on a new plate every 24 hours for a total of 2 transfers. The brood size of each worm was calculated by counting the number of embryos and larvae laid on the three plates. Embryonic lethality was measured by counting the number of the unhatched embryo (dead) 24 hours post laying compared to total embryos laid.

### Counting of oocytes in population

For the WT (N2) and CSR-1 catalytic mutant, germlines of adult worms (72 h post-hatching) were dissected and stained with DAPI and number of oocytes were counted.

### Sorting

Large populations of Late L4 larvae stage from the synchronised population were sorted using the COPAS BIOSORT instrument (Union Biometrica), according to manufacturer’s guidelines. The population was sorted using two size parameters, Time Of Flight (TOF) and Extinction. Stage of the sorted population was validated by counting worms under a microscope by scoring features like closed vulva and absence of oocytes as a characteristic of late L4 stage larvae. First-generation homozygotes for CSR-1 KO or CSR-1 ADH were sorted by excluding GFP positive heterozygote worms. *klp-7_co* heterozygote strain was sorted using GFP marker and GFP positive worms were sorted.

### Immunostaining

Immunostaining was performed as described previously (Barucci et al., 2020). Primary antibody, anti-Flag (Sigma, F1804) at a dilution of 1:500, was incubated overnight at 4°C in PBS, 0.1 % Tween-20, 5 % BSA. The secondary antibody, anti-mouse (Invitrogen, Alexa Fluor 488) at a dilution of 1:500 was incubated for 1 hour at room temperature. DNA was stained with DAPI.

### live imaging

Transgenic worms were mounted on 2% agarose pads in the presence of 0.5% sodium azide. Images were acquired on ZEISS LSM 700 microscope with a ×40 objective or ×63 objective for the mCherry-CSR-1, mCherry-GLH-1 and PIWI-GFP. Images were acquired using the ZEISS ZEN software and processed using ImageJ v.2.0.0.

### Western blotting

Worms were lysed in 1X NuPAGE LDS sample buffer (ThermoFisher Scientific) and heated at 90°C for 10 minutes. Any debris was removed by centrifuging at 18000×g. ∼50 µg of protein extracts was then resolved on precast NuPAGE Novex 4–12% Bis-Tris gels (Invitrogen, NP0321BOX). The proteins were transferred to a nylon membrane with the semidry transfer Pierce Power System (ThermoFisher Scientific) using the preprogrammed method for high-molecular-mass protein. The primary antibodies used included anti-KLP-7 ((Gerson-Gurwitz et al., 2016); a gift from the Desai laboratory) and anti-tubulin (Ab6160, Abcam) antibodies, and the secondary antibodies used included anti-rabbit (31460, Pierce), anti-mouse (31430, Pierce) and anti-rat (A9037, Sigma) HPR antibodies. The SuperSignal West Pico PLUS Chemiluminescent Substrate was used to detect the signal using a ChemiDoc MP imaging system (Biorad).

### RNA extraction

For total RNA extraction, synchronous and sorted populations of ∼1000 worms as described for individual experiments were frozen in dry ice with TRIzol™ (Invitrogen, Ref. 15596026). After five repetitions of freeze and thaw, total RNA was isolated according to the manufacturer’s instructions. For RNA extraction after IP, TRI Reagent was directly added to beads, and RNA extraction was performed as per manufacturer’s instructions. For RNA used for RNA-seq or RT-qPCR, DNase treatment was performed using a maximum of 10 µg RNA treated with 2 U Turbo DNase (Ambion) at 37 °C for 30 min followed by acid phenol extraction and ethanol precipitation. An Agilent 2200 TapeStation System was used to evaluate the RIN indexes of all of the RNA preps, and only samples with RNA integrity number (RIN) > s8 were used for downstream applications.

### RT-qPCR

Reverse transcription was performed according to manufacturer’s instructions using M-MLV reverse transcriptase (Invitrogen, Ref. 28025013) and qPCR was performed using Applied Biosystems Power up SYBR Green PCR Master mix following the manufacturer’s instructions and using an Applied Biosystems QuantStudio 3 Real-Time PCR System. Primers used for qPCR are listed in Table S6.

### IP/ total-sRNA-seq

Total RNA from at least 1000 sorted worms with RIN>9 was used to generate small RNA libraries. For 22G-RNAs from IP, IP was performed using ∼10,000 synchronized and sorted worms for FLAG-CSR-1 or ∼70,000 for GFP-HRDE-1. Worms were lysed in small RNA IP buffer (50 mM HEPES pH 7.5, 500 mM NaCl, 5 mM MgCl2, 1 % NP-40, 10 % glycerol, 1x Halt protease inhibitors and RNaseIn 40 U/mL), using a chilled metal dounce. Crude lysates were cleared of debris by centrifuging at 18,000×g at 4°C for 10 minutes. 10 % of the extract was saved as input, and total RNA was extracted using TRIzol™ as described above. Thre rest of the extract was incubated with 30 µl of Anti-FLAG M2 Magnetic Agarose Beads suspension (Sigma M8823) or 25 µl GFP-Trap Magnetic Agarose (Chromotek) for FLAG-CSR-1 or GFP-HRDE-1 respectively, for 1 hour at 4°C. After four washes of the beads with the small RNA IP buffer, the RNA bound to the bait was extracted by adding TRIzol™ to beads as described above. The library preparation was performed essentially as described previously (Barucci et al., 2020). Amplified libraries were multiplexed to purify further using PippinPrep DNA size selection with 3% gel cassettes and the following parameters for the selection: BP start (115); BP end (165). The purified libraries were quantified using the Qubit Fluorometer High Sensitivity dsDNA assay kit (Thermo Fisher Scientific, Q32851) and sequenced on a NextSeq-500 Illumina platform using the NextSeq 500/550 High Output v2 kit 75 cycles (FC-404-2005).

### Gro-seq

One thousand synchronised and sorted Late L4 worms for WT (N2), *csr-1* catalytic mutant and *csr-1* KO were collected as described above. Nuclear Run-on reaction was performed by incorporating 1 mM Bio-11-UTP, followed by RNA extraction and biotinylated nascent RNA enrichment as described previously (Quarato et al., 2020). Libraries were prepared by repairing 5’-OH of fragmented RNAs by Polynucleotide Kinase (Thermo scientific), followed by 3’and 5’adaptor ligation as described previously (Quarato et al., 2020). Adaptor ligated RNA was reverse transcribed using SuperScript IV Reverse Transcriptase (Thermo Fisher Scientific) following manufacturer conditions except that reaction was incubated for 1h at 50°C. cDNA was PCR amplified with specific primers using Phusion High fidelity PCR master mix 2x (New England Biolab) for 18-20 cycles. Libraries were analysed on Agilent 2200 TapeStation System using high sensitivity D1000 screentapes and quantified using the Qubit Fluorometer High Sensitivity dsDNA assay kit (Thermo Fisher Scientific, Q32851). Multiplexed libraries were sequenced on a NextSeq-500 Illumina platform using the NextSeq 500/550 High Output v2 kit 75 cycles (FC-404–2005).

### Strand-specific RNA-seq library preparation

DNase-treated total RNA with RIN > 8 was used to prepare strand-specific RNA libraries. Ribosomal and mitochondrial rRNAs were depleted using a custom RNAse-H-based method to degrade rRNAs using complementary oligos as described previously (Barucci et al., 2020).

Strand-specific RNA libraries were prepared using at least 100 ng of rRNA depleted RNAs using NEBNext Ultra II Directional RNA Library Prep Kit for Illumina (E7760S). RNA libraries were analysed on Agilent 2200 TapeStation System using high sensitivity D1000 screentapes and quantified using the Qubit Fluorometer High Sensitivity dsDNA assay kit (Thermo Fisher Scientific, Q32851). Multiplexed libraries were sequenced on a NextSeq-500 Illumina platform using the NextSeq 500/550 High Output v2 kit 75 cycles (FC-404– 2005).

### Ribo-seq

Ribo-seq has been performed as described in (Aeschimann et al., 2015) with some modifications (Quarato et al., 2020). Briefly, 10,000 late L4 worms were sorted using COPAS biosorter as described above and were lysed by freeze grinding in liquid nitrogen in Polysome buffer (20 mM Tris-HCl pH 8, 140 mM KCl, 5 mM MgCl2, 1 % Triton X-100, 0.1 mg/mL cycloheximide) and ∼1 mg extract was digested by RNase I (100 U) at 37°C for 5 min. Debris was clarified by centrifuging at 18,000 ×g followed by fractionation on a discontinuous sucrose gradient (10-50 %) by ultracentrifugation at 39,000 rpm for 3 h in an SW41-Ti rotor (Beckman coulter). Monosome fractions were collected by pumping of Fluorinert FC-40 and using a fraction collector by measuring UV trace. RNA extracted from the monosome fraction was DNase treated as described above and fragments of 28-30 nucleotides were size selected by resolving on a 15 % TBE-Urea gel. 3’phosphate was removed (PNK buffer pH 6.5 (70 mM Tris pH 6.5, 10 mM MgCl2, 1mM DTT), T4 PNK (Thermo Scientific), RNaseIN 40 U/mL, 20 % PEG400) and 5’ end was phosphorylated by treating RNA with T4 Polynucleotide Kinase (1x PNK buffer (Thermo Scientific), 1mM ATP). 28-30 nucleotide Ribosome protected fragments (RPF) were then cloned with the sRNA-seq library preparation approach, as described previously (Barucci et al., 2020; Quarato et al., 2020).

### IP-MS/MS

IPs for the MS/MS analysis were performed as described previously (Barucci et.al., 2020). Briefly, Synchronous population of 120,000 (for CSR-1 IPs for RNase treatment or control condition) worms were harvested at 48 hours post-hatching or 20,000 (for CSR-1 IPs comparing WT IP with catalytic mutant) worms were harvested and sorted at 44 hours post hatching and lysed by using a chilled metal dounce in the IP buffer (50 mM HEPES pH7.5, 300 mM NaCl, 5 mM MgCl2, 10% Glycerol, 0.25% NP40, protease inhibitor cocktails (Fermentas). Crude lysates were cleared of debris by centrifuging at 18,000 g at 4°C for 10 minutes. For RNase treatment, RNase I (Invitrogen) 50 U/mg of extract was used at 37 °C for 5 minutes. ∼5 mg of protein extract (for CSR-1 IPs in RNase or control condition) or 1 mg of protein extract (for CSR-1 IPs comparing WT IP with catalytic mutant) was incubated with 15 µl of packed Anti-FLAG M2 Magnetic Agarose Beads (Sigma M8823) for 1 hour at 4°C. After four washes with the IP buffer, the beads were washed twice with 100 µL of 25 mM NH4HCO3. Finally, beads were resuspended in 100 µL of 25 mM NH4HCO3 and digested by adding 0.2 µg of trypsin/LysC (Promega) for 1 h at 37 °C. Samples were then loaded into a homemade C18 Stage Tips for desalting (principally, by stacking one 3M Empore SPE Extraction Disk Octadecyl (C18) and beads from SepPak C18 CartridgeWaters into a 200 µl micropipette tip). Peptides were eluted using a ratio of 40:60 MeCN: H2O + 0.1% formic acid and vacuum concentrated to dryness. Peptides were reconstituted in injection buffer (2:98 MeCN: H2O + 0.3% TFA) before nano-LC-MS/MS analysis as described previously (Barucci et.al., 2020).

## Data analysis

### Sequencing data analyses

Analysis for RNA-seq, sRNA-seq, and GRO-seq have been performed as previously described (Barucci et al., 2020; Quarato et al., 2020). Unless otherwise stated, computations were done using Python and UNIX utilities, either as standalone scripts or as steps implemented in a Snakemake (Koster and Rahmann, 2012) workflow. The scripts and workflows are available at https://gitlab.pasteur.fr/bli/bioinfo_utils. For Ribo-seq data (data analysis pipeline available at the same address), the analysis was performed according to the following steps. The 3′ adapter was trimmed from raw reads using Cutadapt v.1.18 (Martin, 2011) using the following parameter: -a TGGAATTCTCGGGTGCCAAGG - discard-untrimmed. The 5’ and 3’ UMIs were removed from the trimmed reads using cutadapt with options -u 4 and -u -4. After removing UMIs, the reads from 28 to 30 nt were selected using bioawk (https://github.com/lh3/bioawk, git commit fd40150b7c557da45e781a999d372abbc634cc21).

The selected 28–30-nucleotide reads were aligned to the *C. elegans* genome sequence (ce11, C. elegans Sequencing Consortium WBcel235, with an added extra chromosome representing the codon-optimized klp-7 for some libraries) using Bowtie2 (Langmead and Salzberg, 2012) v.2.3.4.3 with the following parameters: -L 6 -i S,1,0.8 -N 0.

Reads mapping on sense orientation on annotated protein-coding genes were considered as Ribosome-protected fragments (RPF). Such reads were extracted from mapping results using samtools (Li et al., 2009) 1.9 and bedtools (Quinlan and Hall, 2010) v2.27.1. RPF reads of size 29 were further classified into subcategories, based on the codons found at the positions corresponding to the A (16-18 nt) and P (13-15 nt) sites of the ribosome. Codon optimality was defined as explained below. Those reads were re-mapped on the genome using bowtie2 (version 2.3.4.3) with options -L 6 -i S,1,0.8 -N 0. The resulting alignments were used to generate bigwig files with a custom bash script using bedtools version v2.27.1, bedops (Neph et al., 2012) version 2.4.35 and bedGraphToBigWig version 4. Read counts in the bigwig file were normalized by million “non-structural” mappers, that is, reads of size 28 to 30 nt mapping on annotation not belonging to the “structural” (tRNA, snRNA, snoRNA, rRNA, ncRNA) categories, and counted using featureCounts (Liao et al., 2013) v.1.6.3. These bigwig files were used to generate “metaprofiles” where normalized coverage information (RPM for Reads Per Million) was averaged across replicates, and represented along sets of selected genes. The metaprofiles were generated using a Python script based on the deepTools (Ramírez et al., 2016) and gffutils (https://github.com/daler/gffutils) libraries. Translation efficiency was calculated as the ratio of TPMs of Ribo-seq and RNA-seq.

### Distance distribution analyses

The distribution of the distances between re-mapped RPF and 22G-RNA-seq reads was computed by counting distances between 5’ ends of RPF and 22G-RNA reads of opposite strandedness, only considering 22G-RNA reads within a distance of +/- 120 bp from the RPF read, and only considering RPF reads mapping in the sense direction within the coordinates of a gene among a selected list. Counts were transformed into z-scores using the Scipy (Virtanen et al., 2020) library (version 1.3.2). A plot of distance distribution, within the (−15, 45) distance range, was made using the Matplotlib library (https://doi.org/10.1109/MCSE.2007.55) version 3.1.1. This was done using z-scores in order to have comparable values between different combinations of libraries. A plot of dominant periods in distance distribution signal was made using the Matplotlib library (https://doi.org/10.1109/MCSE.2007.55) version 3.1.1. The dominant periods were obtained using the fast Fourier transform function of the Scipy library (version 1.3.2) (Virtanen et al., 2020). This was done using z-scores in order to have comparable values between different combinations of libraries.

### Analysis of codon usage

All protein-coding genes were categorized based on their Translation Efficiency in the following categories Log2TE ≥3, ≥2, ≥1, ≤ -1, ≤ -2 and ≤ -3. Relative synonymous codon usage was calculated for genes in each category using the CAI calculator (Puigbò et al., 2008). To calculate enrichment of codons usage in each of the categories, differential RSCU of respective categories of genes was calculated by normalizing their RSCU with RSCU of genes showing a TE of ∼1 (Log2TE 0±0.1). Codons enriched in highly translated mRNAs (Log2TE ≥ 3) were considered optimal codons and codons that were avoided were considered non-optimal. Similarly, differential RSCU analysis was performed for CSR-1 targets.

### Gene ontology

Gene ontology was performed using WormCat tool (Holdorf et al., 2019).

### tRNA copy number and TPM

tRNA copy number was determined using tRNAscan-SE (Chan and Lowe, 2019). TPMs for the tRNAs were extracted from the GRO-seq dataset from WT late L4 staged worms.

### Determination of a codon-optimized sequence for klp-7

The codon-optimized sequence for *klp-7* was computed with a Python script using BioPython (Cock et al., 2009) as follows: To each amino acid, a corresponding optimal codon was associated based on a given optimality ranking. Here, the codon ranking was based on usage in highly translation efficient proteins as explained above. Then, each codon in the CDS of the native *klp-7* gene was replaced with the optimal codon associated to the corresponding amino-acid. For mapping purposes, the resulting sequence was added to the genome as if it was an extra chromosome, and the transgene was added to the annotation files used for read counting. In order to produce comparable bigwig tracks between libraries obtained on different strains (codon-optimized or not), a Python script based on the pyBigWig library (https://doi.org/10.5281/zenodo.594045) was used to relocate the values on the extra chromosome to the actual genomic position of *klp-7. This script is available at https://gitlab.pasteur.fr/bli/libhts*.

### MS/MS Data analysis

For identification, the data were searched against the C. elegans (CAEEL) UP000001940 database (Taxonomy 6239 containing one protein sequence par gene) using Sequest HT through Proteome Discoverer (v.2.2). Enzyme specificity was set to trypsin, and a maximum of two missed cleavage sites was allowed. Oxidized methionine and N-terminal acetylation were set as variable modifications. Maximum allowed mass deviation was set to 10 ppm for monoisotopic precursor ions and 0.6 Da for MS/MS peaks. The resulting files were further processed using myProMS v.3.9 (Poullet et al., 2007) (work in progress). False-discovery rate (FDR) was calculated using Percolator and was set to 1% at the peptide level for the whole study. Label-free quantification was performed using peptide extracted ion chromatograms (XICs), computed with MassChroQ v.2.2.1 (Valot et al., 2011). For protein quantification, XICs from proteotypic peptides shared between compared conditions (TopN matching) with missed cleavages were used. Median and scale normalization was applied on the total signal to correct the XICs for each biological replicate (N=4). To estimate the significance of the change in protein abundance, a linear model (adjusted on peptides and biological replicates) based on two-tailed T-tests was performed, and p-values were adjusted using the Benjamini–Hochberg FDR. Proteins with at least 3 total peptides in all replicates, a 2-fold enrichment and an adjusted p-value < 0.05 were considered significantly enriched in sample comparisons. The MS proteomics data have been deposited to the ProteomeXchange Consortium via the PRIDE (Vizcaíno et al., 2016) partner repository with the dataset identifier PXD012557 and PXD020293.

### Statistics and reproducibility

Almost all the experiments shown in this study were performed independently at least twice, and no inconsistent results were observed. IP and MS experiments were conducted with four biological replicates. Ribo-seq was performed using three biological replicates. All the RNA-seq experiments, GRO-seq, sRNA-seq, IP-sRNA-seq, were performed using two biological replicates. RT-qPCRs to test RNAi efficiency in samples for sequencing experiments were performed in their respective biological experiments. RT-qPCRs for gene expression changes otherwise was performed with at least three biological replicates. Most of the graphs were generated using GraphPad Prism 8. For details of the particular statistical analyses used, precise P-values, statistical significance and sample sizes for all of the graphs, see the figure legends.

### Gene lists

The gene lists used are provided in Table S1.

## DATA AND MATERIALS AVAILABILITY

All sequencing data (GRO-seq, RNA-seq, and sRNA-seq from total lysate or IP experiments, Ribo-seq) are available at the Gene Expression Omnibus (GEO) under accession code <u>GSE155077</u>. The MS proteomics data have been deposited to the ProteomeXchange Consortium via the PRIDE partner repository with the dataset identifier <u>PXD012557</u> and PXD020293. All other data supporting the findings of this study are available from the corresponding author on request. The custom scripts generated for this study are available from the corresponding author on request. Some of the data analysis pipelines are available at https://gitlab.pasteur.fr/bli/bioinfo_utils

## SUPPLEMENTAL FIGURES, TITLES AND LEGENDS

**Figure S1.**
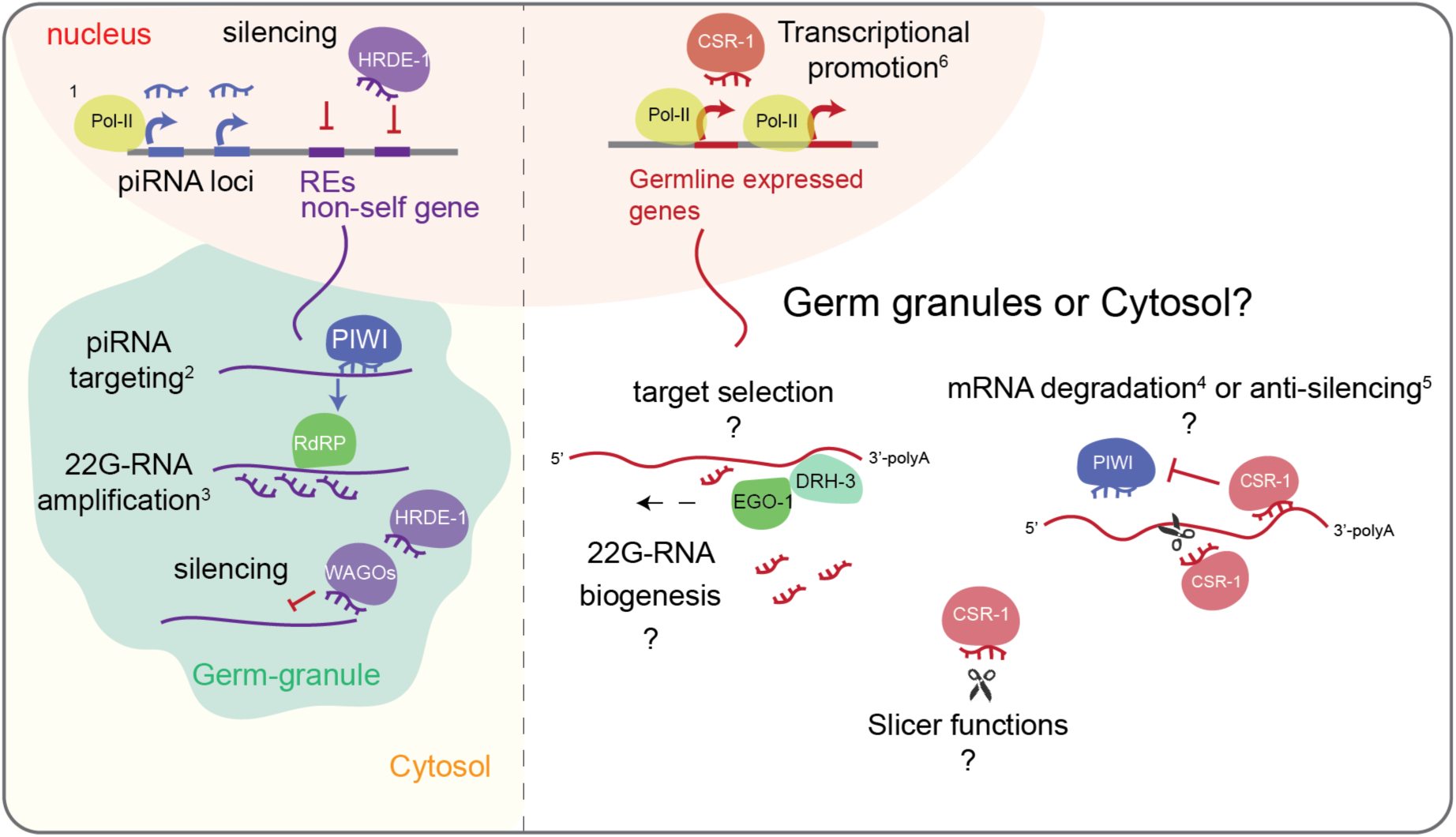
Current model for germline endogenous 22G-RNAs biogenesis and functions. PIWI interacting-RNAs (piRNAs, 21U-RNAs) are transcribed by RNA Polymerase II^1^(Cecere et al., 2012; Gu et al., 2009). piRNAs are then loaded by PIWI which recruit an RNA-dependent RNA polymerase in germ granules to produce secondary antisense small RNAs (22G-RNAs) loaded by WAGOs including nuclear Argonaute HRDE-1 and silence foreign transcripts like REs^2,3^(Bagijn et al., 2012; Batista et al., 2008; Das et al., 2008; Lee et al., 2012). CSR-1 loads 22G-RNAs synthesized by RdRP EGO-1 (Claycomb et al., 2009). No trigger for CSR-1 22G-RNAs is known. Also, it is not clear where CSR-1 22G-RNAs are produced. CSR-1 possess a catalytic activity demonstrated in vitro (Aoki et al., 2007). There is still a lack of clarity on different proposed functions of CSR-1 and its slicer activity. Slicer activity is proposed to degrade mRNA either for maternal mRNA clearance in embryo^4^(Quarato et al., 2020) or fine-tuning of mRNA levels to be delivered to oocytes^4^(Gerson-Gurwitz et al., 2016). On the other hand, it is also proposed as an anti-silencer to PIWI mediated silencing of germline genes mRNAs^5^(Seth et al., 2013; Wedeles et al., 2013) and promote transcription of targets^6^(Cecere et al., 2014). However, it is not clear how CSR-1 targets are selected and how the RdRP EGO-1 mediated biogenesis of 22G-RNAs happen and where these activities take place-germ granule or cytosol?

**Figure S2.**
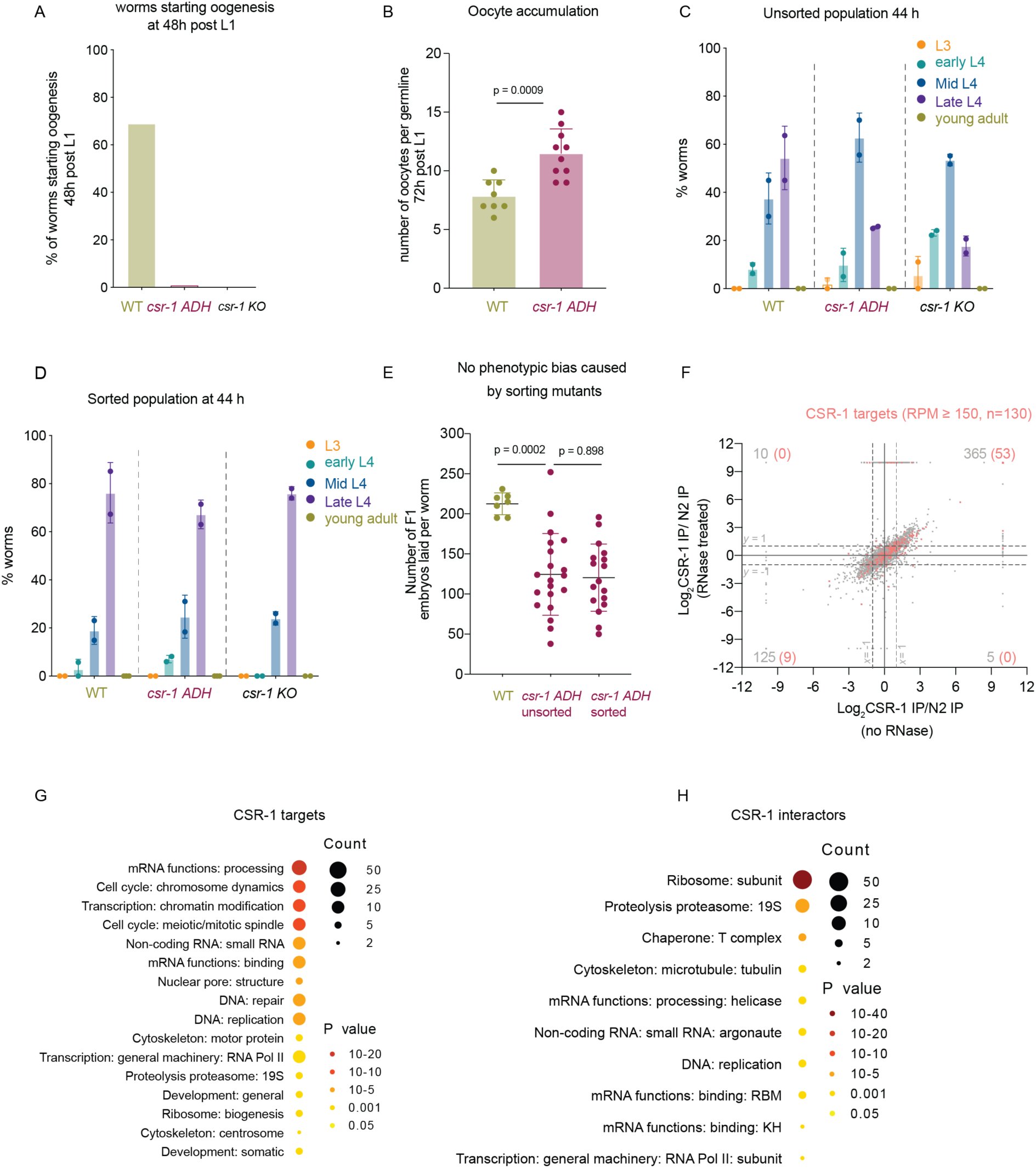
related to Figure 1. Phenotypic characterization of CSR-1 mutants. Comparison of the number of worms with oocytes at 48 h between WT strain, CSR-1 ADH and CSR-1 KO (n = 60 worms). **(B)** Comparison of the number of oocytes present in the germline of adult worms at 72 h post-hatching between WT strain and CSR-1 ADH (n = 10 worms). **(C)** Distribution of WT, CSR-1 ADH and CSR-1 KO population in different larval stages after synchronization at 44 h post-hatching. **(D)** Distribution of WT, CSR-1 ADH and CSR-1 KO population in different larval stages after sorting the synchronized worms at 44 h post-hatching. **(E)** Brood size of CSR-1 ADH strain before and after sorting. Data are represented as mean ± s.d. Two-tailed P values were calculated using Mann–Whitney–Wilcoxon tests. **(F)** Scatter plot comparing the log2 fold changes in CSR-1 interactors (IP-MS/MS) to control IPs performed in WT strain in the absence of RNase treatment (Barucci et al., 2020) (x-axis) to the IPs performed after RNase treatment (this study). CSR-1 targets with 22G-RNA ≥150 RPM are highlighted. Number in grey refers to all interactors with log2 fold change of ≥1 and p-value ≤ 0.05 for each quadrant. The number in parenthesis is CSR-1 targets with 22G-RNA ≥ 150 RPM. Enrichment factor for CSR-1 targets is 6.1 with p < 1.6e^-28^. n = 4 biologically independent experiments. **(G, H)** gene categories enriched in CSR-1 targets (22G-RNA ≥ 150 RPM) **(G)** or CSR-1 interactors **(H)**. Figures generated using WormCat (Holdorf et al., 2019)

**Figure S3.**
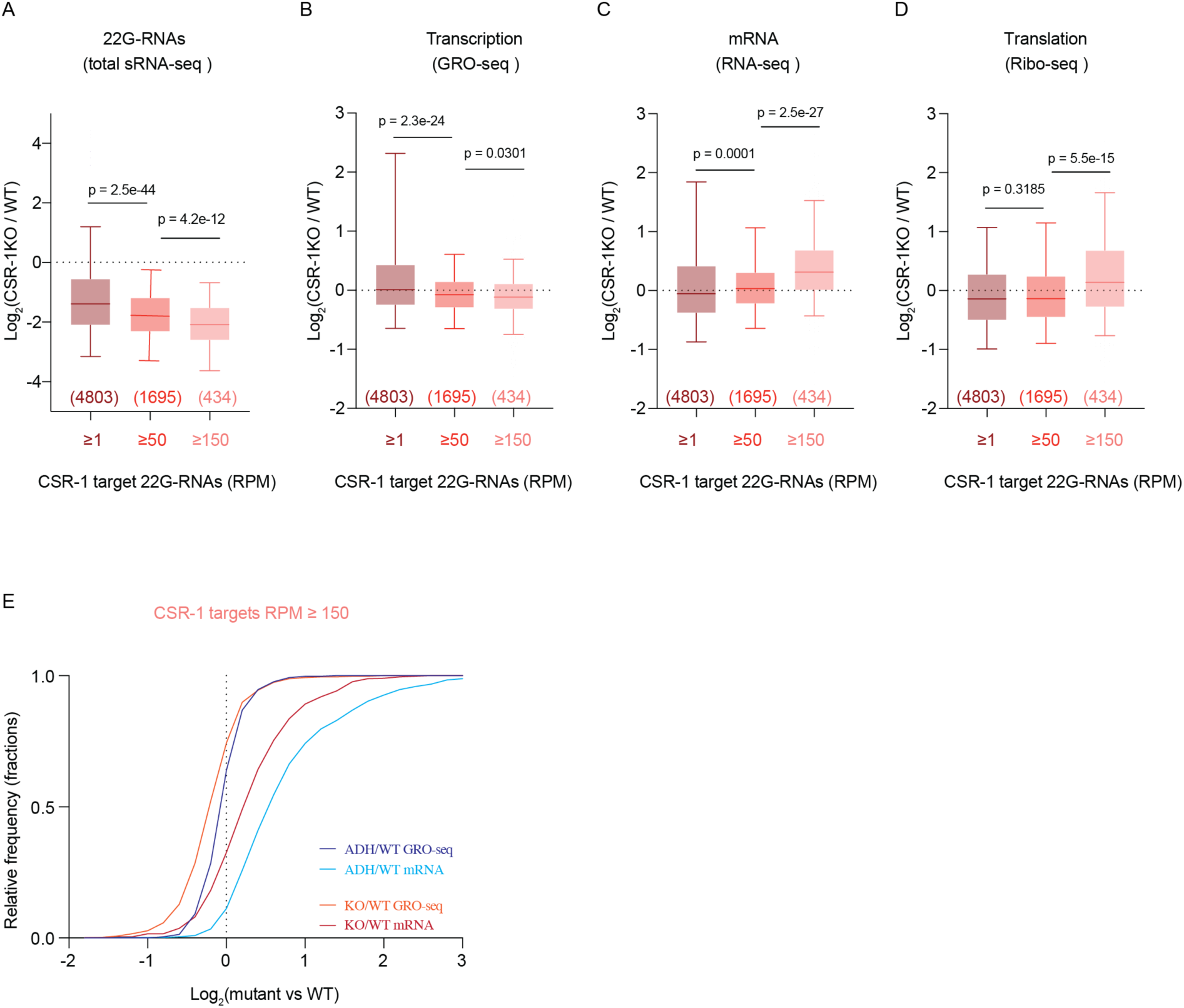
related to Figure 1. Gene expression analyses in *csr-1* KO mutant. **(A-D)** Box plots showing the log2 fold change of total 22G-RNAs (sRNA-seq) (n = 1) **(A);** nascent RNAs (GRO-seq) (**B**); or mRNAs (RNA-seq) **(C);** or mRNAs engaged in translation (Ribo-seq) (**D**), in CSR-1 KO compared to WT strain. The distribution for the CSR-1 targets with 22G-RNA in CSR-1 IP ≥ 1 RPM, ≥ 50 RPM, or ≥ 150 RPM is shown. Box plots display median (line), first and third quartiles (box), and 5^th^ /95^th^ percentile value (whiskers). n = 2 biologically independent experiments. **(E)** Cumulative frequency distribution for CSR-1 targets with RPM ≥ 150. The comparison shows GRO-seq (p = 2e^-11^) and RNA-seq (p = 9e^-19^) for CSR-1 KO or CSR-1 ADH compared to WT. Two-tailed P values were calculated using Mann–Whitney–Wilcoxon tests.

**Figure S4.**
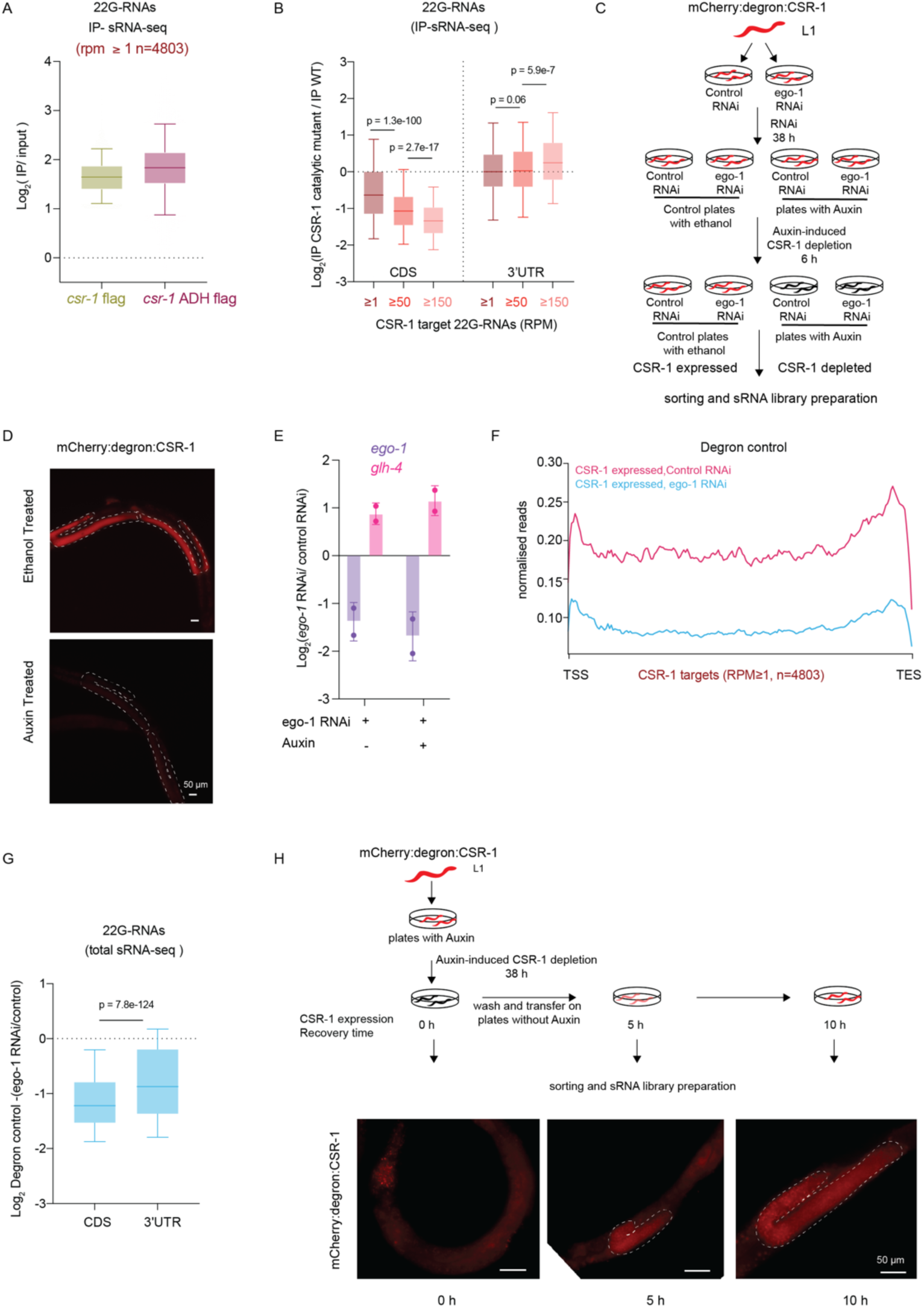
related to Figure 2. CSR-1 catalytic activity regulates the biogenesis of 22G-RNAs. **(A)** Box plot showing log2 fold change in 22G-RNAs in IPs of CSR-1 ADH and to WT CSR-1 IPs compared to input for CSR-1 targets. Box plots display median (line), first and third quartiles (box), and 5^th^ /95^th^ percentile value (whiskers). Two-tailed P values were calculated using Mann–Whitney–Wilcoxon tests, n = 2 biologically independent experiments. **(B)** Box plot showing log2 fold change in 22G-RNAs in IPs of CSR-1 ADH compared to WT CSR-1 on CDS and 3’UTR. The distribution for the CSR-1 targets with 22G-RNA in CSR-1 IP ≥ 1 RPM, ≥ 50 RPM, or ≥ 150 RPM is shown. Box plots display median (line), first and third quartiles (box), and 5^th^ /95^th^ percentile value (whiskers). Two-tailed P values were calculated using Mann–Whitney–Wilcoxon tests, n = 2 biologically independent experiments. **(C)** Experimental scheme and **(D)** fluorescent images of live animals expressing Degron::mCherry::3xFLAG::HA:: CSR-1 used for CSR-1 depletion experiments for Figures 2 D-F, and S4 D-G. Auxin treatment completely depletes CSR-1 in the germline. **(E)** Log2 fold change in expression of RdRP *ego-1* mRNA and *glh-4* (CSR-1 target misregulated upon CSR-1 depletion) upon *ego-1* RNAi compared to control RNAi by qPCR for two biologically independent replicates used for sequencing 22G-RNAs in Figure 2 D-F and S4 D-G. Data shows mean ± s.d. **(F)** Metaprofile analysis as in (Figure 2D) showing the distribution of normalized total 22G-RNA reads (RPM) across CSR-1 targets (22G-RNA ≥1 RPM) upon *ego-1* RNAi and Control RNAi treated in degron control. **(G)** Box-plot as in (Figure 2F) showing the log2 fold change in the amount of 22G-RNA generated from CDS and 3’ UTR of CSR-1 targets (22G-RNA ≥1 RPM) in *ego-1* RNAi compared to control RNAi treated in degron control. **(H)** Experimental scheme and fluorescent images of live animals expressing Degron::mCherry::3xFLAG::HA::CSR-1 used for CSR-1 expression recovery for 0, 5 or 10 h after depletion for 38 h experiments for Figure 2G, H.

**Figure S5.**
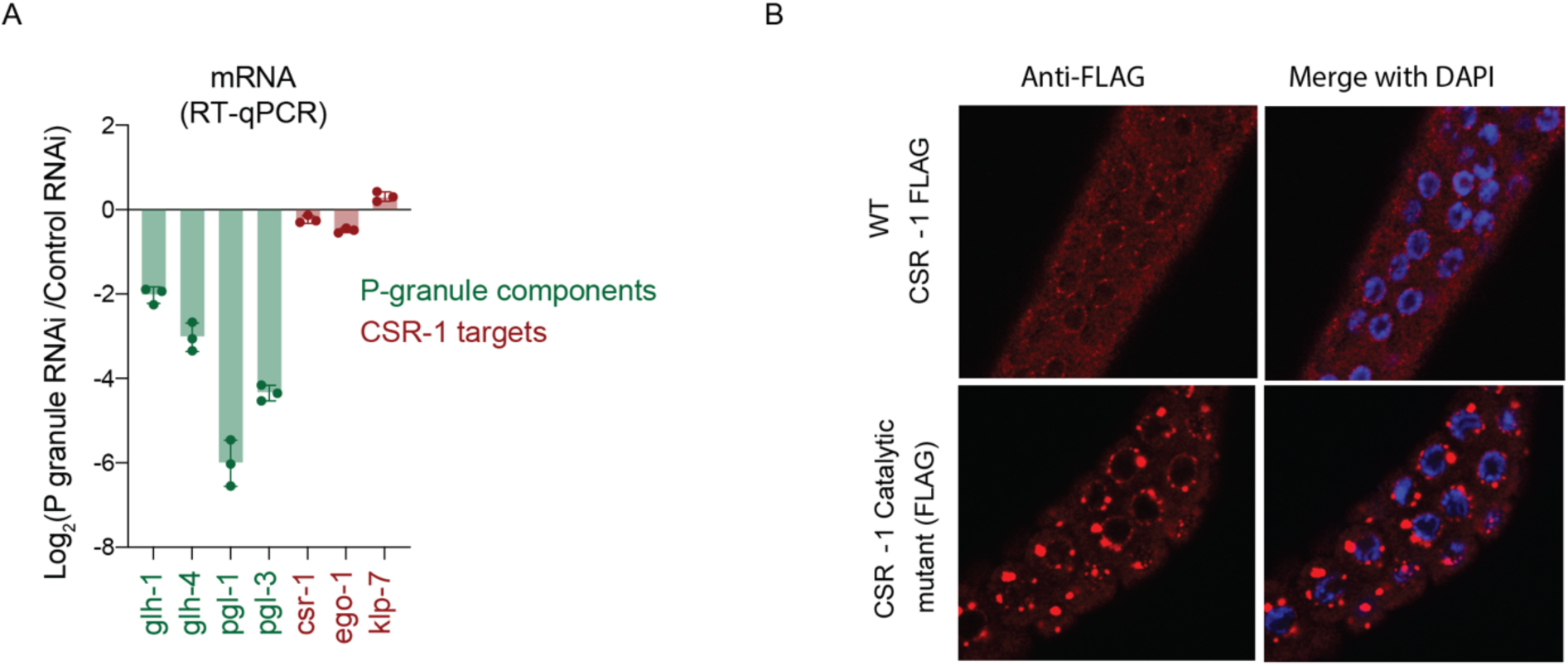
related to Figure 3 and 4. Removal of P granule components does not upregulate CSR-1 targets. **(A)** log2 fold change in expression of *glh-1, glh-4, pgl-1* and *pgl-3* (germ granule components) and *csr-1, ego-1* and *klp-7* (CSR-1 targets) upon P granule RNAi compared to control RNAi by RT-qPCR for the two biologically independent replicates for sRNA-seq in Figure 3B and C. **(B)** Immunofluorescence showing localization and expression of FLAG-tagged WT CSR-1 and CSR-1 ADH and a merge with DAPI stained nucleus. In WT, CSR-1 is localized to the cytosol and P granule. In CSR-1 catalytic mutant, CSR-1 is predominantly localized in enlarged P granules (Brightness of catalytic mutant decreased to avoid image saturation due to higher expression levels of mutant protein).

**Figure S6.**
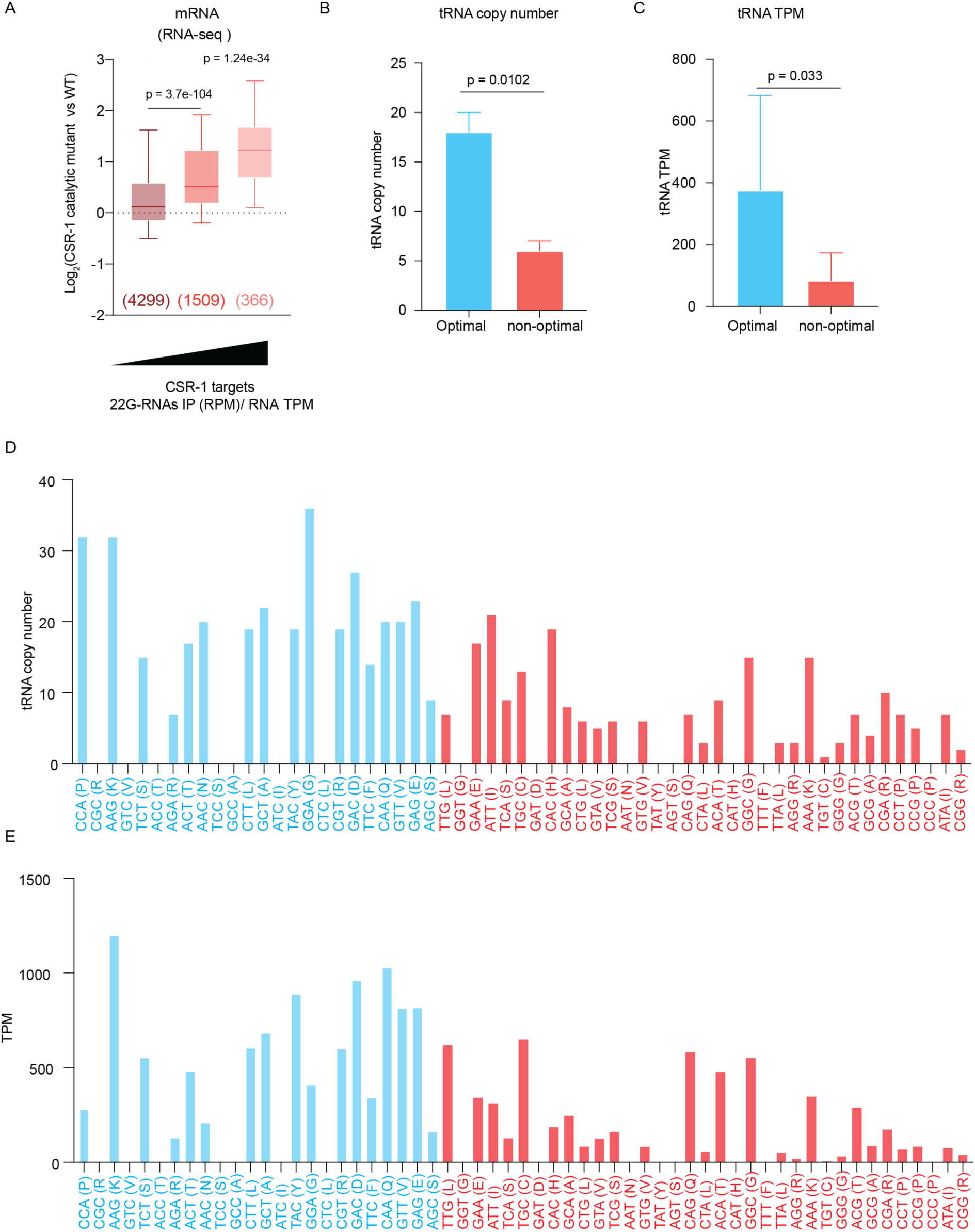
related to Figure 5. tRNA copy number and expression for optimal and non-optimal codons. **(A)** Box plots showing the log2 fold change of mRNAs (RNA-seq) in CSR-1 ADH compared to WT strain (same as Figure 1D) except, CSR-1 targets are ranked based on 22G-RNA density by normalizing 22G-RNA reads by RNA TPMs to account for RNA abundance. **(B)** Bar graph showing the median copy numbers for tRNAs for optimal or non-optimal codons with a 95 % confidence interval as in Figure 5C without adjusting for absent tRNAs, **(C)** Bar graph showing the median TPMs for tRNAs from the GRO-seq dataset for WT strain at the late l4 larval stage (44h) for optimal or non-optimal codons with a 95 % confidence interval as in Figure 5D, without adjusting for absent tRNAs. Two-tailed P values were calculated using Mann–Whitney–Wilcoxon tests, n = 2 biologically independent experiments. **(D)** Plot showing the copy number of tRNAs in the genome for each codon. **(E)** Plot showing the availability of the tRNA pool corresponding to each codon (TPM for tRNAs from GRO-seq, n = 2 biologically independent experiments).

**Figure S7.**
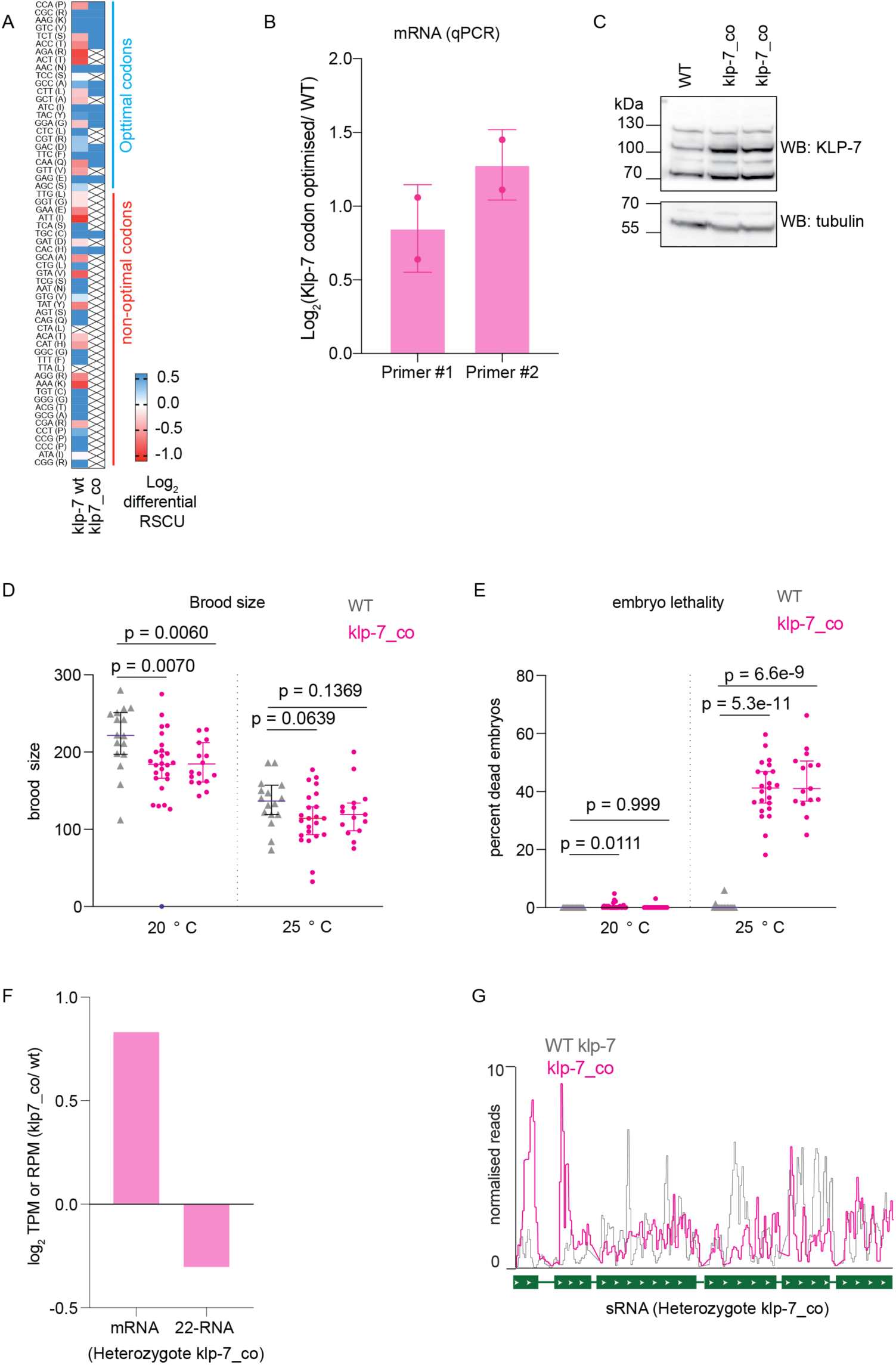
related to Figure 6. Effects of codon optimization of CSR-1 target - *klp-7*. **(A)** heat map showing log2 fold change in RSCU for either WT *klp-7* or *klp-7_co* compared to genes showing neutral translational efficiency of 1 as explained in methods (similar to Figure 5B, C). The blue line highlights optimal codons in genes with high TE, and the red line highlights non-optimal codons. × denotes absence of the codon in the coding sequence. **(B)** log2 fold change in expression of *klp-7* in the *klp-7* codon-optimized strain, *klp-7_co*, compared to WT *klp-7* in WT strain by RT-qPCR for replicates used for sequencing in Figure 6. **(C)** Immunoblot showing expression of the KLP-7 protein in WT strain and two independent CRISPR-Cas9 lines where endogenous *klp-7* was replaced by modified *klp-7_co*, immunoblot for alpha-tubulin serves as the loading control. **(D)** Brood size of WT strain and two independent CRISPR-Cas9 lines where modified *klp-7_co* replaced endogenous *klp-7* at 20° and 25° C. **(E)** Embryonic lethality observed in WT strain and the two independent CRISPR-Cas9 lines where modified *klp-7_co* replaced endogenous *klp-7* at 20° and 25° C. Data are represented as mean ± s.d. Two-tailed P values were calculated using Mann–Whitney–Wilcoxon tests. **(F)** Plot showing log2 fold change for normalized reads for mRNA and 22G-RNAs for the codon-optimized copy of *klp-7_co* compared to WT copy *klp-7* from heterozygote worms, n=1. **(G)** A genomic view of normalized reads of 22G-RNA reads antisense to WT copy of *klp-7* (Grey) and codon-optimized *klp-7_co* copy (purple) from heterozygote worms.

## SUPPLEMENTAL DATA TABLES

**Table S1:** Gene lists generated and used in this study

**Table S2:** IP-MS/MS - CSR-1 IP / control IP

**Table S3:** IP-MS/MS - CSR-1 ADH IP / CSR-1 WT IP

**Table S4:** List of strains

**Table S5:** List of guide and repair template sequences used to generate CRISPR-Cas9 alleles

**Table S6:** Oligo pairs used for RT-qPCR

